# NicheSphere reveals Spp1⁺ macrophages as central hubs coordinating fibrotic remodeling in myeloproliferative neoplasms

**DOI:** 10.64898/2026.03.16.711605

**Authors:** Hélène F.E. Gleitz, Mayra L. Ruiz Tejada Segura, James S. Nagai, Stijn N.R. Fuchs, Gerjanne Vroeg in de wei, Inge A.M. Snoeren, Giulia Cesaro, Iris J. Bakker, Marta Gargallo Garasa, Jessica E. Pritchard, Tesa Klenovšek, Stephani Schmitz, Lina Schmidt, Eric Bindels, Twan Lammers, Joost Gribnau, Hind Medyouf, Kai Markus Schneider, Carolin V. Schneider, Rafael Kramann, Ivan G. Costa, Rebekka K. Schneider

**Affiliations:** Department of Developmental Biology, Erasmus Medical Center, Rotterdam, the Netherlands; Oncode Institute, Erasmus Medical Center Cancer Institute, Rotterdam, the Netherlands; Department of Cell and Tumor Biology, Faculty of Medicine, University Hospital RWTH Aachen, Aachen, Germany; Institute for Computational Genomics, Faculty of Medicine, RWTH Aachen University, Aachen, Germany; Center for Computational Life Sciences, RWTH Aachen University, Aachen, Germany; Department of Information Engineering, University of Padova, Italy; Department of Hematology, Erasmus Medical Center Cancer Institute, Rotterdam, the Netherlands; Institute for Experimental Molecular Imaging, RWTH Aachen University, Germany; Department of Oncology, Hematology and Stem Cell Transplantation (Medical Clinic IV); Center for Regenerative Therapies Dresden (CRTD), TUD Dresden University of Technology, Dresden, Germany; Department of Medicine 1, University Hospital Carl Gustav Carus Dresden, Technische Universität (TU), Dresden, Germany; Else Kroener Fresenius Center for Digital Health, Medical Faculty Carl Gustav Carus, TUD Dresden University of Technology, Dresden, Germany; Department of Medicine III, University Hospital RWTH Aachen, Aachen, Germany; Institute of Experimental Medicine and Systems Biology, RWTH Aachen University, Medical Faculty, Aachen, Germany; Division of Nephrology and Clinical Immunology, RWTH Aachen University, Medical Faculty, Aachen, Germany; Department of Internal Medicine, Nephrology and Transplantation, Erasmus Medical Center, Rotterdam, The Netherlands

## Abstract

Bone marrow fibrosis in myeloproliferative neoplasms arises from interactions between mutant hematopoietic clones and fibrosis-driving stromal cells. We identify Spp1⁺ macrophages as central “communication hubs” integrating inflammatory and fibrotic programs *via* spatial proximity, ECM signaling, and cytokine activation. Using dual lineage-tracing, single-cell and multiplet RNA-sequencing, and a novel computational method for cell-colocalization and communication analysis named NicheSphere, we show that Spp1⁺ macrophages form core communication hubs with osteoCAR cells, fibroblasts, and megakaryocytes. NicheSphere uncovered two distinct niches: macrophage-enriched compartments driving WNT, JAK-STAT, and TNFα cytokine signaling, and a fibrosis-interacting core enriched in TGF-β and ECM glycoproteins. Genetic ablation revealed cooperative roles of stromal and hematopoietic Spp1 in sustaining fibrosis and inflammation. Mechanistically, SPP1 promoted integrin-mediated adhesion, IL-1β secretion, and stromal activation, while IL-1 cytokines induced Spp1 and collagen expression. Loss of Spp1 in hematopoietic progenitors reduced inflammation and restored macrophage function, establishing SPP1⁺ macrophages as therapeutic targets in progressive bone marrow fibrosis.

## Introduction

Myeloproliferative neoplasms (MPN) are initiated by the acquisition of somatic mutations in hematopoietic stem cells (HSCs) that lead to clonal expansion and abnormal blood cell development. The majority of patients harbor mutations in the Janus kinase signal transducer JAK2 (JAK2^V617F^), the chaperone protein calreticulin (CALR) or the thrombopoietin receptor (MPL). Primary myelofibrosis (PMF) is the most aggressive MPN, characterized by progressive bone marrow fibrosis, aberrant megakaryocyte proliferation, cytopenias, constitutional symptoms, and extramedullary hematopoiesis.

PMF is driven not only by MPN driver mutations but also by the aberrant secretion of chemokines and cytokines that remodel the bone marrow (BM) niche. Among these mediators, the chemokine platelet factor-4 (PF4, also known as CXCL4) is markedly up-regulated in MF and elevated PF4 levels correlate with disease stage, promote myofibroblast differentiation and extracellular matrix (ECM) deposition of (Gli1+) fibrosis-driving mesenchymal stromal cells (MSCs), and amplify pro-inflammatory JAK/STAT signaling in both megakaryocytes and stromal populations^1,2^. Additionally, receptor-ligand interactions in human PMF bone marrow compared to controls suggested a megakaryocyte-driven PF4 cross-talk with fibrosis driving mesenchymal stromal cells (MSCs) in PMF^1,3^. However, while megakaryocyte-derived PF4 was shown to reprogram (Gli1+) MSCs and drive their fibrotic remodeling, emerging evidence suggests that additional PF4-secreting hematopoietic subsets may similarly engage—and be engaged by—the stromal niche through both soluble and contact-dependent mechanisms.

Recent advances in single-cell RNA-sequencing (scRNA-seq) and high-resolution computational methods have begun to resolve the cellular heterogeneity and intercellular interactions that underlie BM fibrosis^4^. Conventional scRNA-seq workflows capture the transcriptomes of individual cells but lose spatial context and direct evidence of physical interactions. Spatial protocols and transcriptomics are revolutionizing our understanding of cellular interactions^5–9^, but the bone marrow is challenging for these applications because the trabecular and cortical bone must be decalcified—often compromising RNA integrity—while the highly heterogeneous, tightly packed marrow niche and abundant extracellular matrix impede uniform sectioning, and accurate spatial resolution of diverse cell–cell interactions.

To overcome these challenges, we have adapted a sequencing of physically interacting cell protocol (PIC-seq^10^) for PMF that preserves doublet information, allowing us to trace all PF4-expressing hematopoietic cells in the bone marrow (tracked *via* Pf4-ZsGreen) and their interaction with Gli1-tdTomato+ fibrosis-driving cells (after transplantation into Gli1;tdTom recipient mice). We have strong morphological evidence for the direct proximity of Gli1+ fibrosis-driving and PF4+ hematopoietic cells. By sorting both singlets and ZsGreen+/tdTomato+ multiplets, we can compare the transcriptional program of these cell populations and their direct cellular interaction. A yet open challenge is the computational analysis of PIC-seq data. Previous work used either custom scripts^10^ or computational tools^11^. However, these methods struggle to quantify how cell–cell interactions change between homeostasis and disease. Likewise, ligand–receptor analyses offer functional insights from single cell data but do not account for direct physical contacts^12^. To bridge this gap, we developed NicheSphere, a computational framework that integrates physical interaction data from PIC-seq with ligand–receptor–based communication inferred from paired single-cell and multiplet/doublet RNA-seq. Moreover, NicheSphere leverages curated ligand-receptor pairs, such as ones related to immune responses, extracellular matrix (ECM) or user-defined sets, to characterize cell communication mechanisms within and between niches. By applying NicheSphere to Pf4-ZsGreen⁺ hematopoietic cells and Gli1-tdTomato⁺ stromal cells in PMF, we identified cellular and molecular niches with increased interactions influencing both ECM remodeling and immune recruitment that drive disease progression.

## Results

### Single-cell sequencing of physically interacting cells in bone marrow fibrosis identifies fibrosis-driving single cells and interacting multiplets

To analyze pro-fibrotic interactions of PF4-expressing hematopoietic cells influencing the fibrotic transformation of the bone marrow (BM) in PMF, we employed a dual lineage tracing approach using Pf4-ZsGreen+ ckit-enriched hematopoietic cells transduced with a TPO-overexpression lentiviral vector (blue fluorescent protein; BFP backbone) or its empty vector (EV) as a control, transplanted into lethally-irradiated bi-genic Gli1CreER;tdTomato mice (Fig. 1A-B, Supplementary Fig. 1A). This allowed us to track the entirety of PF4-expressing cells, specifically megakaryocytes, monocytes, macrophages, and granulocytes (Supplementary Fig. 1B) as well as Gli1^+^tdTomato^+^ stromal cells that we have identified as fibrosis-driving cells^2^. TPO overexpression, which models MPN and BM fibrosis by signaling through the MPL receptor and robustly activating the JAK-STAT signaling pathway, induced significant myeloproliferative features including leukocytosis, thrombocytosis and splenomegaly (Supplementary Fig. 1B-D, 1F). The frequency of Pf4-ZsGreen+ hematopoietic cells significantly increased over time confirming the TPO-induced myeloproliferation. Confocal imaging confirmed the increase of both Pf4+ hematopoietic cells and Gli1+ stromal cells in the TPO model of MPN and MF in sub-endosteal and perivascular localization (Fig. 1A). Importantly, accumulations of Pf4+Gli1+ cells were found in perisinusoidal localization. In contrast, in the EV condition, spindle-shaped Gli1+ cells line the endosteal bone and Pf4+ cells are mostly positioned in perisinusoidal locations.

**Figure 1.**
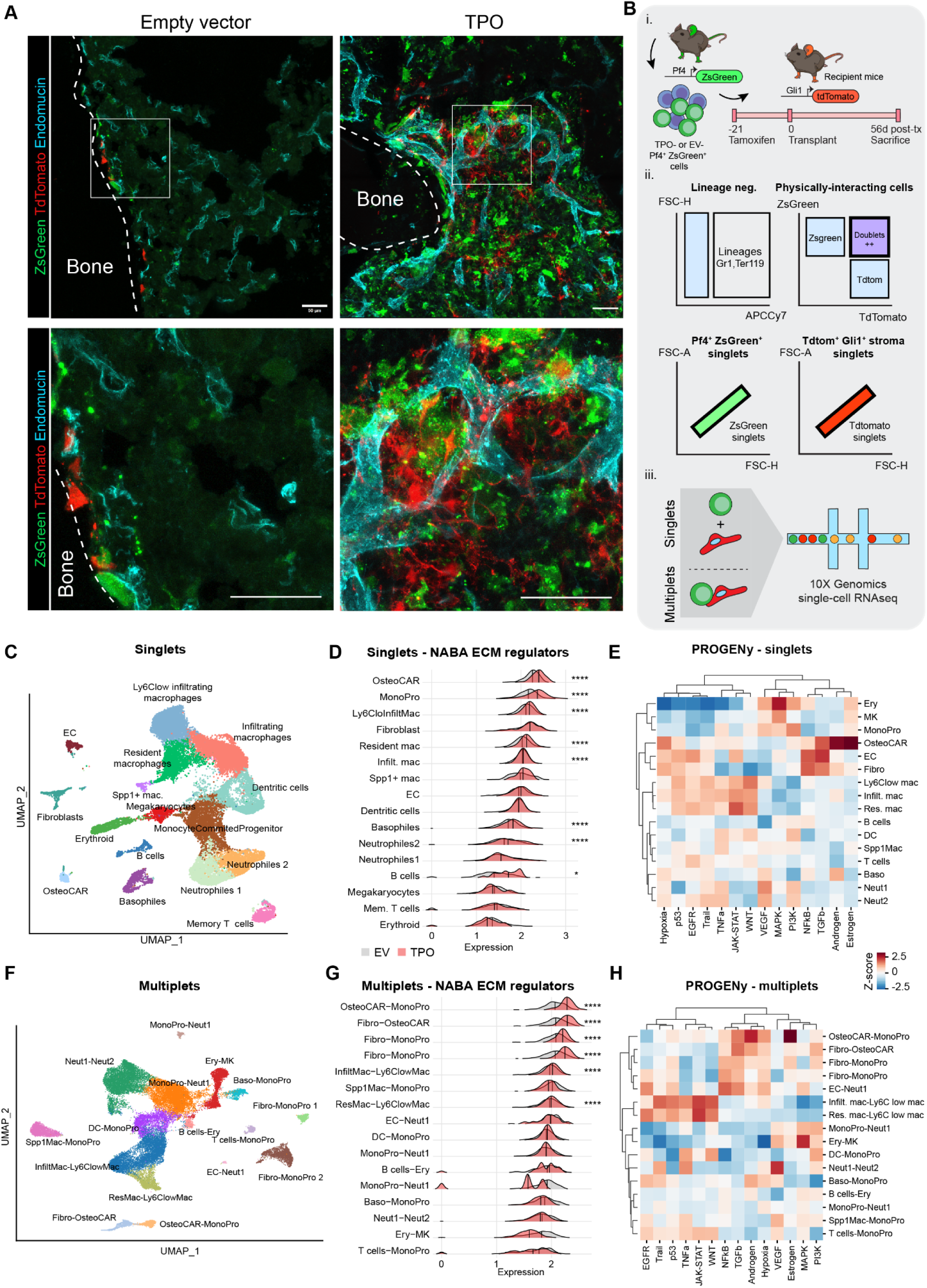
Physically Interacting Cell sequencing (PICseq) in TPO-induced primary myelofibrosis. (A) Confocal imaging of Pf4-ZsGreen (green) cells transplanted into Gli1-tdTomato+ (red) recipient mice, overexpressing either TPO or the empty vector (BFP), co-stained with endomucin for vasculature (cyan). Expanded confocal images in lower panels, showing Pf4-ZsGreen (green) cells transplanted into Gli1-tdTomato+ (red) recipient mice and interacting in close proximity to vasculature. Scale bar: 50um. (B) Panel i) shows the schematic representation of the biological experimental design, including transplanting PF4-ZsGreen ckit+ cells transduced with TPO-BFP or EV-BFP into Gli-TdTomato recipient mice. Panel ii) highlights the FACS sorting strategy of ZsGreen+ and tdTomato+ singlets and ZsGreen+TdTomato+ doublets, which are then processed for single-cell RNAsequencing using the 10X Genomics platform in panel iii. (C) UMAP displaying single cells (singlets) sorted for ZsGreen and tdTomato from TPO and EV control mice and sequenced using the 10X Genomics pipeline, n=27,046 singlets. (D) Ridge plot with the composite expression of ECM regulator gene set for singlet clusters. P-value is based on a Wilcoxon test.(*) = p<0.05,(**)= p<0.01, (***) = p<0.001, (****) = p<0.0001. (E) Progeny analysis on singlet clusters. Heatmap of z-score normalized pathway activity scores per cell type. (F) UMAP displaying clusters of multiplet cells sorted for double-positive ZsGreen and tdTomato from TPO and EV control mice and sequenced using the 10X Genomics pipeline, n=35,927 multiplets. (G) Ridge plot with the composite expression of ECM regulator gene set for multiplet clusters. P-value is based on a Wilcoxon test. (*) = p<0.05,(**)= p<0.01, (***) = p<0.001, (****) = p<0.000. (H) Progeny analysis on multiplet clusters. Heatmap of z-score normalized pathway activity scores per multiplet cluster.

To isolate both the Pf4+ and Gli1+ cells separately, but also the identified double-positive cells as physically interacting cells (PICs, Supplementary fig. 2E), we established a gentle dissociation protocol of the long bones to isolate and sort Gli1-tdTomato+ stromal cells, PF4-ZsGreen+ hematopoietic cells and cells that stained double positive for both tdTomato and ZsGreen. The sorting strategy included the isolation of double-positive multiplets and also singlets from each tdTomato or ZsGreen cell populations (Fig. 1B). Two mice per TPO or control group were pooled per sample, and 10,000 ZsGreen+ cells and 60,000 Gli1-tdTomato+ stromal cells were loaded together on one chip, as well as 60,000 double-positive multiplets as a separate sample.

Single cell sequencing recovered 27,046 high-quality singlets across 16 clusters (Fig. 1C; Supplementary Fig. 1G-H). Unsupervised cluster analysis specifically revealed a high granularity of mononuclear phagocytes including monocyte-committed progenitors, infiltrating and donor-derived resident-like macrophages, Spp1+ macrophages and dendritic cells. The recovery of granulocytes as well as CXCL12-abundant reticular (CAR) cells and fibroblasts, further confirmed the successful enrichment of Pf4+ and Gli1+ populations (Supplementary Fig. 1H). Additionally, B-cells, memory T– cells, basophils, endothelial cells and erythroid cells were recovered. Compositional analysis indicated a significant increase in the proportion of infiltrating macrophages and basophils in TPO-induced fibrosis (Supplementary Fig. 1I). OsteoCAR populations showed significantly increased expression of ECM-related genes in TPO-induced fibrosis (Fig. 1D), supporting that they are fibrosis-driving cells. DE gene analysis confirmed increased Tpo expression in all diseased hematopoietic cells but also osteoCAR cells and increased expression of the Tpo ligand and Mpl receptor in megakaryocytes and osteoCAR cells (Supplementary fig. 1J).

By comparing the differential expression signature of singlets with pathway signatures (pathway responsive genes [PROGENy] analysis^13^, we demonstrated that the non-hematopoietic cell types such as endothelial cells, fibroblasts and osteoCARs specifically upregulated TGFβ, the master switch of fibrosis, and NFκB in TPO-induced fibrosis, both pathways known to be involved in the fibrotic differentiation of stromal cells. Erythroid cells and megakaryocytes specifically upregulated MAPK and resident-like and infiltrating macrophages, but not Spp1+ macrophages, upregulated Wnt, Jak-Stat, Tnfa, EGFR and p53 related signaling pathways and thus are central pathways involved in myeloproliferation and inflammation in MPN (Fig. 1E).

We isolated 35,927 multiplets, and based on singlet expression signatures, identified 16 multiplet clusters (Fig. 1E). First, to confirm our dual lineage tracing approach, we asked if multiplets indeed contain both hematopoietic and non-hematopoietic markers. We thus contrasted the number of Pf4-expressing and Gli1-expressing cells in both singlet and multiplet datasets. This analysis indicated a larger number of double positive cells in all multiplet clusters and a small frequency of double positive cells in the singlet clusters (Supplementary Fig. 1L). Next, we applied Bayesprism^14^ to deconvolute the multiplets, and assigned cluster labels based on the two most prevalent cell types within each cluster (Supplementary Fig. 1K). Many multiplet clusters were predominantly composed of monocytes and macrophages, reflecting their abundance among the singlet populations (Fig. 1F). Stromal cells such as fibroblasts and osteoCAR cells were often co-localized with monocyte progenitors and we also observed pure hematopoietic co-localizations as neutrophil-neutrophil or fibroblasts-osteoCAR associations, respectively. In line with the singlet data, doublets containing osteoCARs and fibroblasts showed significant up-regulation of ECM regulators (Fig. 1G). Pathway analysis shows an increase of TGFβ and NFκB signaling in all multiplet clusters with at least one stromal cell type in TPO-induced MF (OsteoCAR-MonoPro, Fibro-OsteoCAR, Fibro-MonoPro, EC-Neutro1) (Fig. 1H). This also mirrored the enrichment of TGFβ and NFκB signaling in non-hematopoietic subsets seen within the singlet data. Infiltrating and resident-like macrophages exhibited upregulation of WNT, JAK-STAT, TNFα, and p53 pathways in TPO-induced fibrosis, suggesting that distinct cell types, even when co-localized with others, are enriched in specific programs—more inflammatory in monocytes and macrophages and more pro-fibrotic in non-hematopoietic stromal cells. However, while this deconvolution approach yielded valuable insights into dominant cell states and signaling pathways, it remained limited in reliably detecting multiplets involving rare but relevant cell types such as megakaryocytes. To overcome this limitation, we sought to develop a new computational approach for the analysis of multiplet data comparing different conditions.

### NicheSphere: A spatial framework to reveal biological communication processes

To tackle the difficulty in the delineation of multiplet cell clusters from rare cells, we devised a new computational approach which we termed NicheSphere. NicheSphere uses the results of a deconvolution algorithm, e.g: BayesPrism^14^, to delineate the identities of all multiplet cells in a condition (disease or healthy) by using the single cell as a reference. This is used to estimate the probability of two cell types interacting in a particular condition. We then used an empirical statistical test to find pairs of cells with significant increase or decrease in interactions in the two conditions (disease vs. healthy). These interactions are encoded as a cell-to-cell graph, where edge weights represent the signed log-odds scores of cellular interactions. NicheSphere only considers pairs with significant interactions changes. Finally, a graph-based clustering reveals niches (or hubs) associated with the conditions (Fig. 2A; Supplementary File 1).

**Figure 2.**
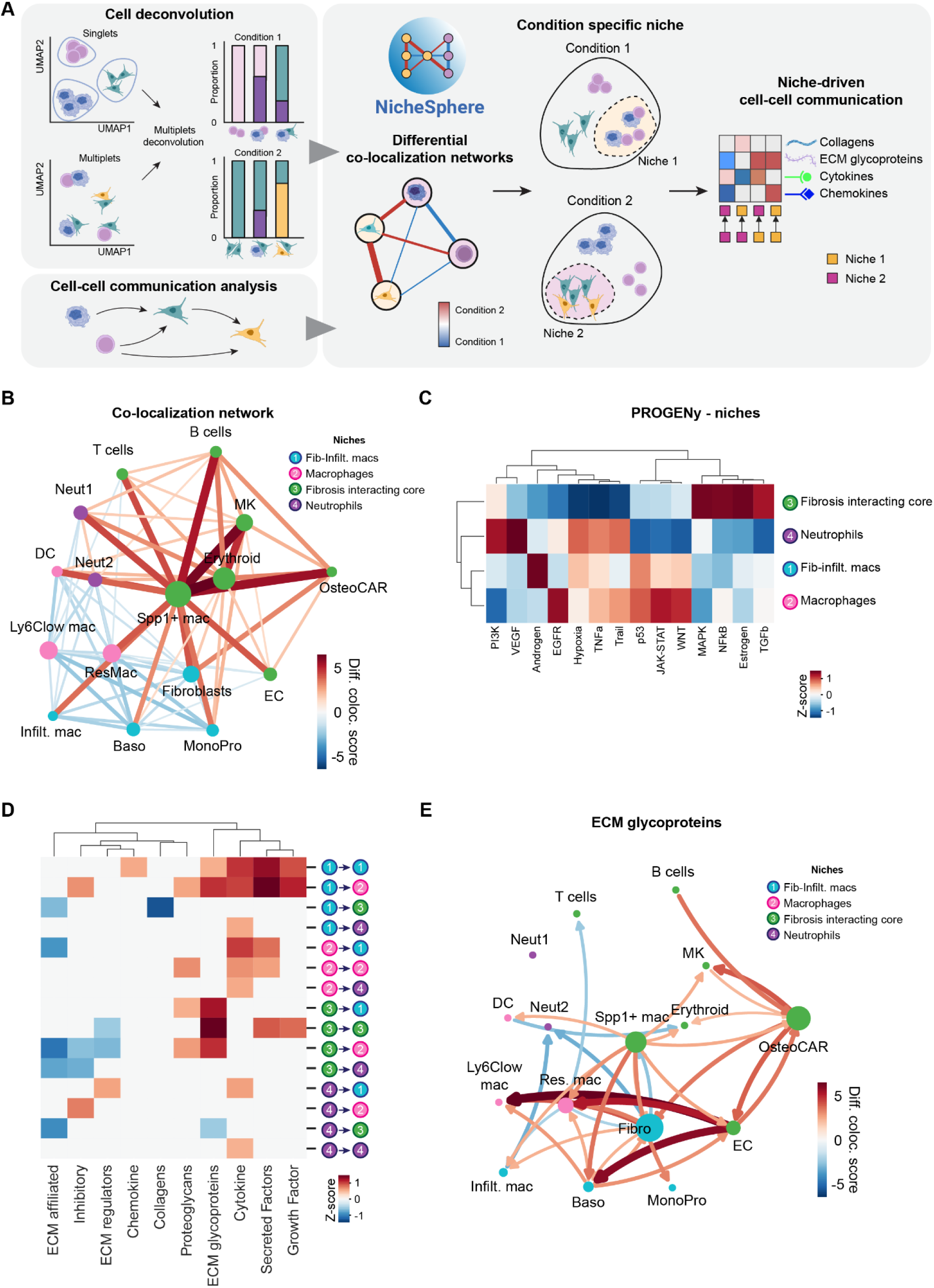
NicheSphere identifies distinct communication programs in spatially co-localized niches. (A) NicheSphere schematics: Using results from deconvolution algorithms in multiplex cells, NicheSphere builds a differential co-localization network and detects groups of cells (Niches) that have an increased or decreased interaction in disease/control conditions. Additionally, using condition specific LR scores obtained via cell a communication tool, NicheSphere can perform biological process based localized differential communication to find biological processes driving changes in cell-colocalization in disease (TPO) or normal conditions. (B) Differential co-localization network showing cell types (nodes) colored by Niche with node size based on betweenness score, i.e. nodes interconnecting other important nodes. Edge colors represent normalized differential co-localization scores for cell type pairs. Red color in edges indicates that cells with increased co-localization in disease (TPO), while blue indicates decrease in colocalization in controls (C) Progeny analysis on singlets grouped by niche. Heatmap of z-score normalized pathway activity scores per niche. (D) Process-based and co-localized differential communication heatmap. Colors represent significant differential communication between differentially co-localized cells. Red indicates increasing signaling in disease (TPO), blue indicates decreasing signaling in controls. Only significant changes (p-values < 0.01; Wilcoxon rank sums test) in cell-cell communication are shown. (E) ECM glycoproteins specific network with cell types (nodes) colored by Niche and node size based on betweenness score. Edge colors represent normalized differential communication scores for cell type pairs. Red color in edges indicates increased signaling in disease (TPO), blue indicates decreased signaling in controls

NicheSphere inferred a co-localization network with 4 communication-hubs: 1) a Fibroblast-Infiltrating macrophage niche (Fib-Infilt. macs) composed of fibroblasts, infiltrating macrophages, basophils and monocyte progenitors), 2) a macrophage niche composed of resident-like macrophages, Ly6C^low^ macrophages and dendritic cells, 3) a niche termed the Fibrosis Interacting Core composed of known, central players in myelofibrosis such as megakaryocytes (MKs) and osteoCAR cells, but also Spp1+ macrophages, erythroid cells, B cells, T cells, and endothelial cells and finally 4) a neutrophil niche with colocalized neutrophil subtypes 1 and 2 (Fig. 2B).

In TPO-induced fibrosis, the Fibrosis Interacting Core showed an increased co-localization score indicating that these cell types have increased interactions in disease (Fig. 2B). Interestingly, Spp1+ macrophages showed an increased co-localization score with almost all other cell types. Additional analyses defining properties to identify cells working as co-localization/interaction hubs, i.e. graph degree, page rank and betweenness, determined Spp1+ macrophages as a central cell in the TPO-specific fibrosis communication hub (Supplementary Fig. 2A). This was particularly interesting as Spp1+ macrophages did not stand out in the previously performed pathway analysis of either singlets or doublets (compare Figs. 1E and 1H). This shows that NicheSphere is able to identify specific roles of cell types in altered cellular interactions in disease and not only differentially expressed genes comparing health and disease. In particular, osteoCAR cells and MKs, both known as central players in MF, had significantly increased co-localization with Spp1+ macrophages. Spp1+ macrophages also showed a significantly increased co-localization with the Fib-infilt. Hub (hub 1) and also the neutrophil hub 4. As Spp1+ macrophages appeared as such a central communication hub, we sought to further characterize this cell population. A more detailed analysis of Spp1+ macrophage markers and signatures indicate them as CD45^+^ CD68^+^ Fabp5^+^ Lpl^+^ Trem2^+^ CD9^+^ cells^15–17^, with a pronounced phagocytic and lipid-associated profile (Supplementary Fig. 2B-E).

Interestingly, the co-localization between hub 1, 2 and 4, which is dominated by mononuclear phagocytes and neutrophils, decreased in TPO-induced fibrosis. Pathway analysis on the different co-localization niches in TPO-induced fibrosis compared to the EV control further delineated the different functions of the different communication niches. Niche 3 (fibrosis interacting core) showed the characteristic up-regulation of the pro-fibrotic TGFβ and NFκB signaling (Fig. 2C). Niche 4 (neutrophils) was enriched in PI3K and VEGF signaling. Niche 1 (Fib-infilt. macs) and niche 2 (macrophages) showed upregulation of the characteristic WNT, JAK-STAT, TNFa and p53 pathways which are characteristic for the fibrosis-inducing hematopoietic clone and also inflammation.

To better understand the cell-cell communication mechanisms underlying the differential co-localization network, NicheSphere includes a curated set of ligand-receptors (LR) related to 10 fibrosis-relevant mechanisms associated with extracellular matrix organization and immune recruitment (Supplementary Table 1; Supplementary Fig. 2F). NicheSphere maps out cell–cell communication by focusing on cell colocalization and constructing ligand–receptor interaction networks^18^ of interacting cell types. It then applies a statistical test to identify which signaling mechanisms are significantly more active—based on the distribution of ligand and receptor expression—within specific niches and under disease conditions.

In TPO-induced fibrosis, the highest increase in cell-cell communication scores were observed between the Fibrosis-Interacting Core niche with nearly all other niches via ECM glycoprotein–mediated communication (Fig. 2D). If we consider only ECM glycoprotein-mediated communication, graph network properties (graph degree) supports the central role of Spp1+ macrophages (Fig. 2E; Supplementary Fig. 2G). OsteoCAR cells emerge as the primary interaction partners in this network, actively engaging with endothelial cells.

The cell communication patterns of different niches are related to distinct fibrosis-related mechanisms (Fig. 2D). The co-localization of the macrophage-enriched niches 1 and 2 showed upregulated cytokines, secreted factors and growth factors, while their interaction with the fibrosis-interacting core was enriched for collagens, proteoglycans and ECM glycoproteins (Fig. 2D, Supp. Fig. 2G). The neutrophil niche in co-localization with the other niches showed a less clear pattern and even upregulation of inhibitory pathways. Altogether, these data indicate that macrophages are recruited to the fibrotic core, including osteoCARs as fibrosis-driving cells, where they contribute to ECM regulation while they promote the inflammatory and secretory processes in bone marrow fibrosis by cytokine and growth factor upregulation when interacting with other mononuclear phagocytes.

### Spp1 marks the fibrosis-interacting core and mediates key ECM glycoprotein interactions

Due to the central role of cellular co-localization in the “fibrosis interacting core” regarding ECM expression, we next interrogated disease-specific ECM glycoprotein interactions in more detail. Visualization of the top disease-specific ligand-receptor interactions related with ECM glycoproteins within the fibrosis-interacting core revealed Spp1 as a ligand associated with top LR pairs in interactions from osteoCAR cells specifically with integrins (e.g. Itgav, Itga9, Itga4, Itgb) as receptors with cells from all other niches (Fig. 3A). Differential expression analysis confirmed the upregulation of Itgav specifically in multiplets containing Spp1+ macrophages, fibroblasts and osteoCAR cells (Supplementary Fig. 3A).

**Figure 3.**
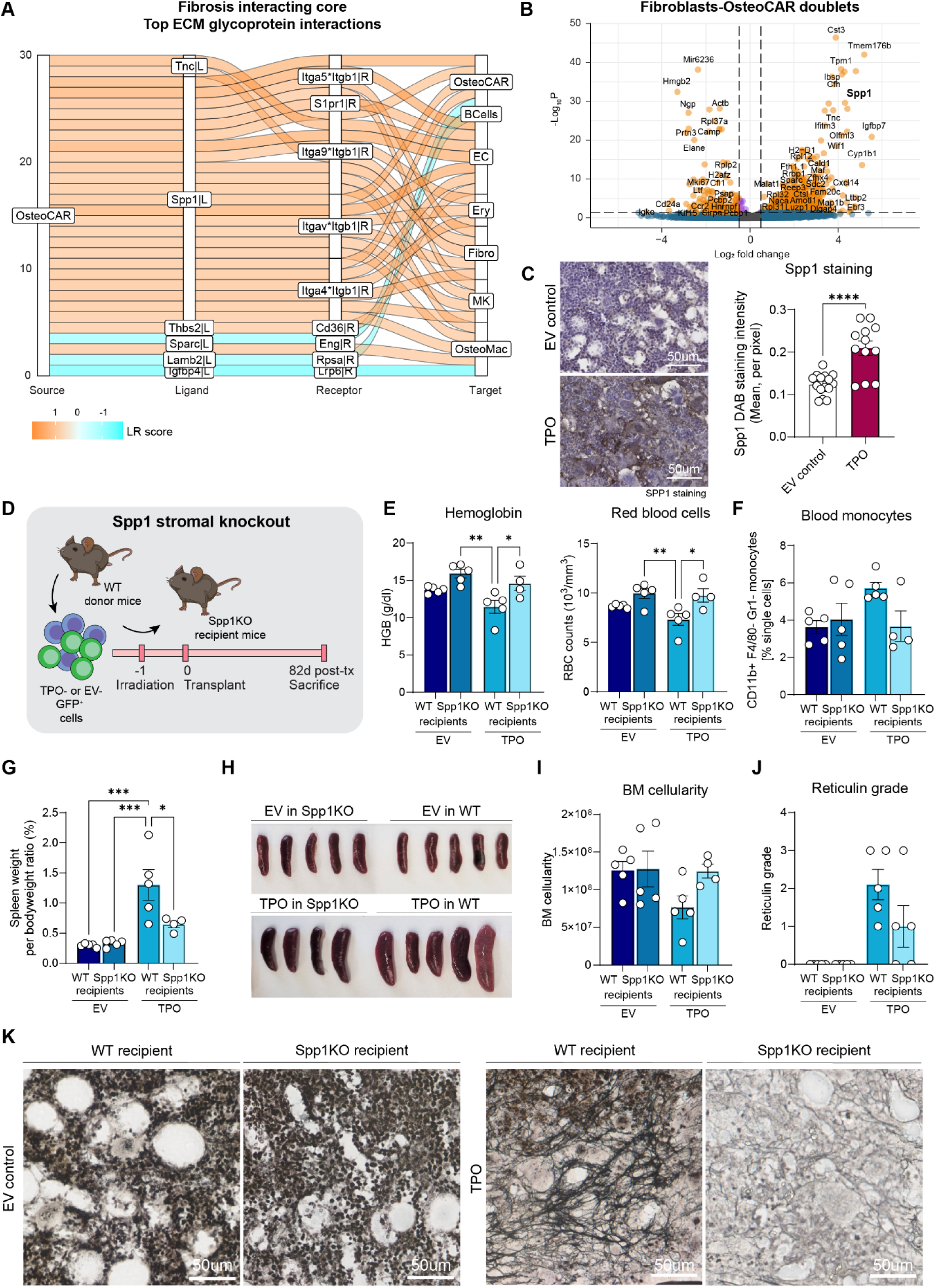
The absence of Spp1 in the stromal compartment ameliorates BM fibrosis and MPN phenotype. (A) Sankey plot showing top 30 differential LR interactions from OsteoCAR cells through ECM glycoproteins towards other co-localized cells. Colors represent LR expression per interaction: orange indicates stronger signaling in disease, blue indicates weaker signaling in disease. (B) Volcano plot for differential gene expression of OsteoCAR-Fibroblast multiplets cell. Genes with logFC > 1 and p-value < 0.05 shown in red; genes with p-value < 0.05 but low logFC in blue and others in green. (C) Representative images of Spp1 staining and quantification of mean DAB intensity in 4um femoral sections of TPO– or EV-induced murine BM fibrosis. Scale bar: 50um. P-value is based on an unpaired t-test.(*) = p<0.05. (D) Schematic representation of experimental timeline, where WT ckit+ donor cells are transduced with TPO-GFP or EV-GFP control lentiviral vectors, and transplanted into lethally-irradiated Spp1KO recipient mice (n=4-5/group). (E) Hemoglobin and red blood cell counts in TPO-driven fibrosis in WT and Spp1KO recipient mice at the time of sacrifice (82 days post-transplant). P-value is based on a one-way ANOVA.(*) = p<0.05,(**)= p<0.01, (***) = p<0.001, (****) = p<0.0001. (F) Blood monocytes quantified by flow cytometry in TPO-driven fibrosis in WT and Spp1KO recipient mice at the time of sacrifice (82 days post-transplant). (G) Spleen weight in TPO-driven fibrosis in WT and Spp1KO recipient mice at the time of sacrifice (82 days post-transplant). P-value is based on a one-way ANOVA.(*) = p<0.05,(**)= p<0.01, (***) = p<0.001, (****) = p<0.0001. (H) Image of spleens isolated from TPO-driven fibrosis in WT and Spp1KO recipient mice at sacrifice. (I) Bone marrow cellularity in TPO-driven fibrosis in WT and Spp1KO recipient mice at the time of sacrifice (82 days post-transplant). (J) Quantification of reticulin staining in 4um femoral sections in TPO-driven fibrosis in WT and Spp1KO recipient mice at the time of sacrifice (82 days post-transplant). (K) Representative imaging of reticulin staining in TPO-driven fibrosis in WT and Spp1KO recipient mice. Scale bar: 50um.

As fibroblasts and osteo-CAR cells showed ECM-enriched co-localization (Fig. 2D) and up-regulation of the TGFβ pathway (Fig. 1H), we specifically interrogated differentially expressed genes in these doublets (Fig. 3B). The significant upregulation of Spp1, also known as osteopontin (OPN), caught our attention. In singlets, the upregulation of Spp1 was specific to osteoCAR cells and Spp1+ macrophages as central cells of the fibrosis-interacting core in TPO-induced bone marrow fibrosis (Supplementary Fig. 3B). In doublets, the expression of Spp1 significantly increased in co-localized cell types containing osteoCAR cells and fibroblasts (Fig. 3B). Spp1 immunohistochemistry and quantification confirmed the increased Spp1 expression in TPO induced bone marrow fibrosis and accentuated in cells with spindle-shaped morphology (Fig. 3C).

As Spp1 was significantly up-regulated in the non-hematopoietic stromal compartment including fibroblasts and osteo-CAR cells specifically with regards to ECM-related cellular interactions, we hypothesized that the stromal knockout of Spp1 would reduce bone marrow fibrosis. To test this, Spp1KO or wild-type (WT) mice were recipients of either TPO-transduced or EV-transduced WT ckit+ hematopoietic stem and progenitor cells (HSPCs; Fig. 3D). In this manner, stromal cells as fibrosis-driving cells do not express Spp1 but HSPCs have normal Spp1 expression. The absence of Spp1 from the stromal compartment normalized the decrease in hemoglobin and red blood cell counts in TPO-induced MF, which are surrogate markers of fibrosis, reflecting the insufficient hematopoiesis in progressed fibrosis (Fig. 3E). The pathognomonic MPN monocytosis was reduced in the absence of Spp1 in stromal cells in TPO-induced fibrosis (Fig. 3F). The increased spleen size as another sensitive marker of progressed fibrosis in TPO-induced fibrosis was significantly reduced in the Spp1 stromal knockout (Fig. 3G-H). Critically, the absence of stromal Spp1 in BM fibrosis restored normal BM cellularity (Fig. 3I) and reduced the reticulin fibrosis grade BM fibrosis (Fig. 3J-K), confirming that stromal Spp1 expression has a central role in bone marrow fibrosis.

### Spp1–IL-1β axis links macrophage signaling to fibroblast inflammatory activation

Having shown that stromal Spp1 expression is crucial to BM fibrosis, we wondered how Spp1+ macrophages as a central co-localization hub in bone marrow fibrosis affect the fibrotic transformation. Spp1+ macrophages, characterized by high expression of Spp1 (osteopontin), have emerged as key pro-fibrotic mediators in several tissues. These macrophages have been implicated in liver, lung, heart and kidney fibrosis, often appearing during disease progression^19–22^. We thus next focused on the receptor-ligand interactions of Spp1+ macrophages. The top ranking ligand was SPP1 communicating to almost all cell types through integrins and CD44 as the receptors (Fig. 4A). We thus identified SPP1 as a key mediator of cellular adhesion and pro-fibrotic crosstalk within the bone marrow microenvironment.

**Figure 4.**
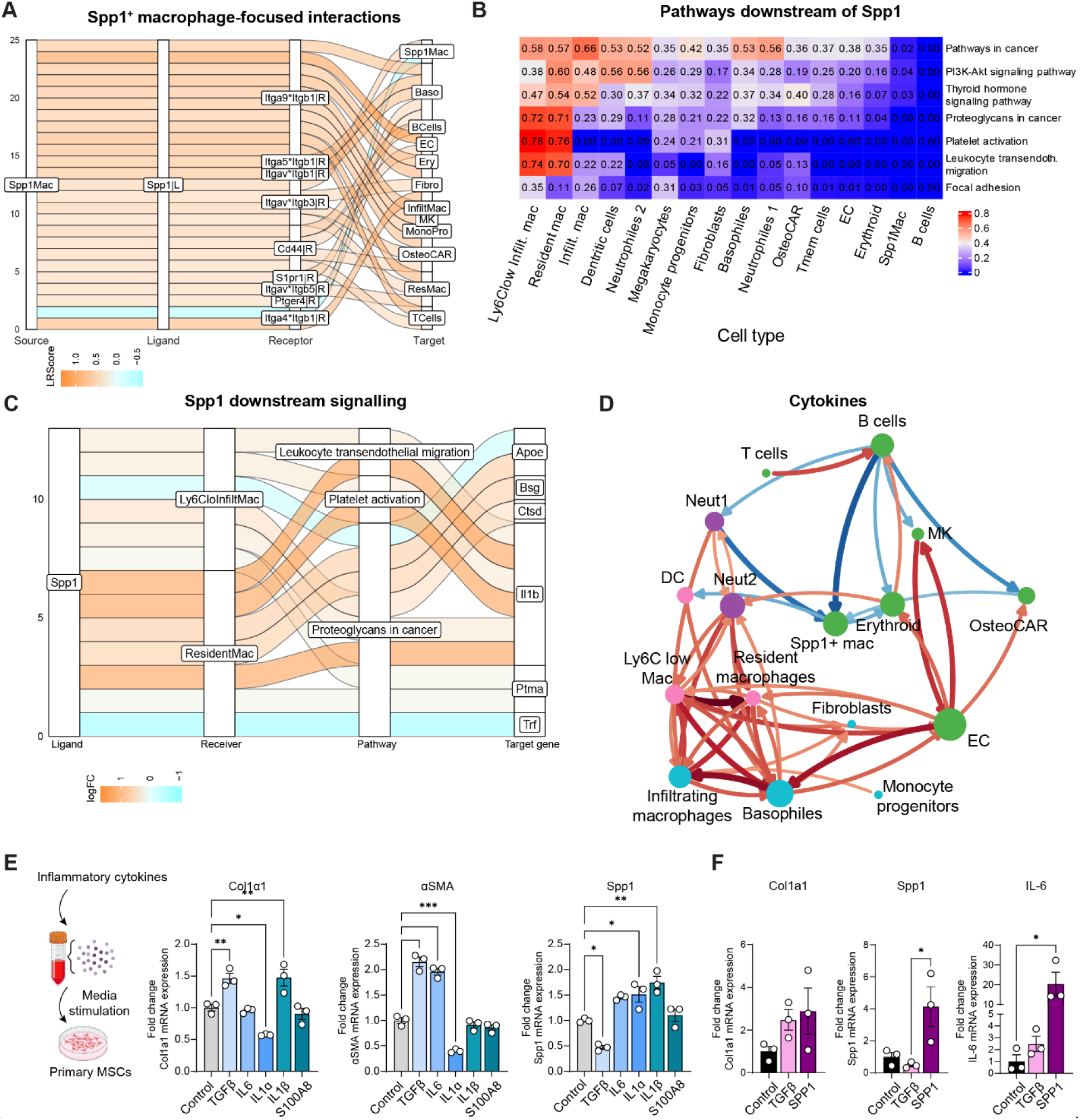
Macrophage-derived Spp1 promotes IL-1β–dependent inflammatory activation. (A) Sankey plot showing top 25 LR pairs associated with Spp1+ macrophage-driven signaling via Spp1 as a ligand. (B) Heatmap of pathway score based on SPP1-associated intracellular signaling, highlighting differential transcriptional response between TPO vs. EV. Higher scores indicate stronger evidence of differential pathway regulation due to up-stream SPP1 signaling. (C) Sankey plot showing SPP1-associated intracellular signaling in Ly6C^low^ infiltrating macrophages and resident-like macrophages, and pathways with enriched differential pathway score from panel B. Genes on the right represent genes downstream of SPP1. Here, only target genes that are ligands are shown. The color map reflects the log-transformed fold change (logFC) between TPO vs. EV. (D) Cytokines localized differential communication network showing cell types (nodes) colored by Niche with node size based on betweenness score. Edge colors represent normalized differential communication scores for cell type pairs. Red color in edges indicates increased signaling in disease, blue indicates decreased signaling in disease. (E) mRNA expression of Col1a1, αSMA and Spp1 in Gli1tdtomato MSCs stimulated with DMSO control, Tgf-β, IL-6, IL-1a, IL-1b and S100A8 for 48 hours, n=3/group, P-value is based on a one-way ANOVA.(*) = p<0.05,(**)= p<0.01, (***) = p<0.001, (****) = p<0.0001. (F) mRNA expression of Col1a1, Spp1 and IL-6 in Gli1tdtomato MSCs stimulated with DMSO control, Tgf-β and recombinant Spp1 for 48 hours, n=3/group, P-value is based on a one-way ANOVA.(*) = p<0.05,(**)= p<0.01, (***) = p<0.001, (****) = p<0.0001.

To understand intercellular interactions downstream of SPP1 better, we next performed an intra-cellular based cell-cell communication analysis^23^. In particular Ly6c^low^ infiltrating macrophages and resident-like macrophages were identified to be activated by SPP1, specifically with regards to KEGG terms as “platelet activation”, “proteoglycans in cancer” and “leukocyte transendothelial migration” (Fig. 4B). This pointed towards SPP1’s role in matrix remodeling and was in line with recent reports showing a role of SPP1 in platelet activation in fibrosis^20^. In addition, we computationally validated for the first time that the intracellular algorithm can recapitulate the bona fide mechanism of MPN in the TPO-induced model of MPN and fibrosis. The analysis further identified an increase in JAK-STAT signaling in megakaryocytes in TPO-induced bone marrow fibrosis and demonstrates that the secretion of TPO from distinct immune cells activates JAK-STAT signaling in megakaryocytes via the MPL receptor (Supp. Fig. 4B).

We next wanted to more specifically understand the downstream effect on Spp1-related pathways (Fig. 4C). This indicated that Spp1 specifically acts on resident-like macrophages, which upregulate the identified pathways leading to increased IL-1β expression as a target gene (Supplementary Fig. 4C-D). We hypothesized that SPP1 activates resident-like macrophages in MPN and fibrosis to acquire a more inflammatory phenotype. We thus specifically interrogated cytokine-driven interactions in the identified niches. Interestingly, while the “fibrosis-driving core” was enriched for “extracellular matrix” (compare Fig. 2), the macrophage-driven niches were enriched for cytokines (Fig. 4D), again highlighting the role of the different niches in fibrosis.

To better understand the effects of pro-inflammatory cytokines such as IL-1β in macrophages on the activation and reprogramming of the stromal compartment, as well as the induction of Spp1, we stimulated primary sort-purified Gli1-tdTomato⁺ stromal cells with pro-inflammatory and pro-fibrotic growth factors and cytokines, including TGF-β, IL-6, IL-1α, and IL-1β, for 48 hours (Fig. 4E). Following stimulation with IL-1α and IL-1β, we observed a significant increase in *Col1a1* and *Spp1* expression, indicating that IL-1 family cytokines robustly promote extracellular matrix production and osteopontin induction. Interestingly, TGF-β treatment led to increased Col1a1 and Acta2 (αSMA) expression, consistent with myofibroblast activation, but resulted in decreased Spp1 mRNA levels. These findings suggest that inflammatory and profibrotic signals engage distinct transcriptional programs in stromal cells, with IL-1-driven pathways preferentially inducing Spp1 as a potential mediator of inflammation-associated niche remodeling in myeloproliferative neoplasms.

### Hematopoietic deletion of Spp1 decreases bone marrow macrophage frequency and reduces IL-1β expression and secretion

Having shown that stromal Spp1 expression is crucial for fibrosis and that Spp1+ macrophages act as central hubs of pro-fibrotic signaling, we next aimed to clarify how hematopoietic-derived Spp1 contributes to bone marrow fibrosis. Our analyses demonstrated that SPP1 is a dominant ligand mediating adhesion and activation across stromal and immune compartments and that it drives resident-like macrophages toward an inflammatory phenotype marked by increased IL-1β expression. These findings suggested that macrophage-derived Spp1 could amplify both matrix remodeling and cytokine-driven stromal activation. To directly test this hypothesis, we generated models with knockout of Spp1 in hematopoietic stem and progenitor cells, including macrophages. We isolated ckit-enriched bone marrow cells from either WT or Spp1KO mice, which were transduced with either TPO-GFP lentiviral vector or its empty vector control. Transduced cells were transplanted into lethally-irradiated WT recipient mice (Fig. 5A). Spp1 knockout in HSPCs in TPO-induced PMF ameliorated the progressive anemia (Fig. 5B) indicative of insufficient hematopoiesis due to bone marrow fibrosis. Flow cytometric analysis revealed a significantly reduced frequency of circulating monocytes in the blood of mice with hematopoietic KO of Spp1 in TPO-induced fibrosis (Fig. 5C). Importantly, reticulin fibrosis was reduced in the absence of Spp1 in hematopoietic cells (Fig. 5D-E). We next asked if the absence of Spp1 in hematopoietic cells would indeed reduce IL-1β downstream of Spp1. IL-1β levels in the blood and also IL-1β positive cells were significantly reduced in the hematopoietic Spp1 knockout (Fig. 5F-G), confirming that hematopoietic Spp1 plays a central role in inflammation and in the consecutive activation of fibrosis-driving cells.

**Figure 5.**
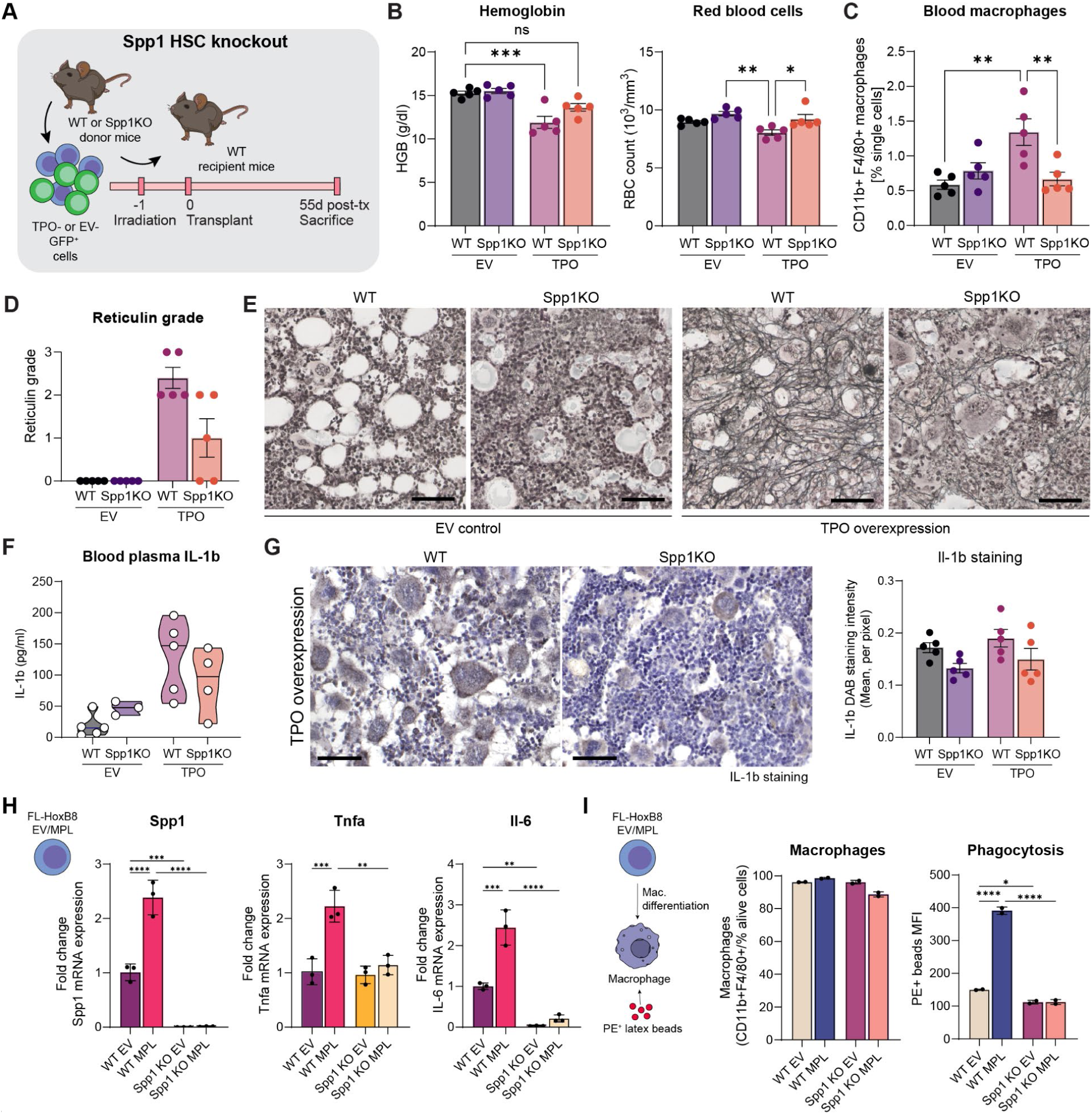
Loss of Spp1 in hematopoietic cells alleviates BM fibrosis and MPN features. A. Schematic representation of experimental timeline, where WT or Spp1KO ckit+ donor cells are transduced with TPO-GFP or EV-GFP control lentiviral vectors, and transplanted into lethally-irradiated WT recipient mice (n=5/group, experiment duration: 55 days after transplant). B. Hemoglobin and red blood cell counts in TPO-driven fibrosis in WT recipient mice with WT or Spp1KO BM at the time of sacrifice (55 days post-transplant). P-value is based on a one-way ANOVA.(*) = p<0.05,(**)= p<0.01, (***) = p<0.001, (****) = p<0.0001. C. Blood CD11b+ F4/80+ monocyte-derived macrophages quantified by flow cytometry in TPO-driven WT or Spp1KO fibrosis in WT recipient mice at the time of sacrifice (55 days post-transplant). D. Quantification of reticulin staining in 4um femoral sections in EV– or TPO-driven fibrosis in WT or Spp1KO-driven fibrosis at the time of sacrifice (55 days post-transplant). E. Representative imaging of reticulin staining in 4um femoral sections in TPO-driven WT or Spp1KO-driven fibrosis in WT recipient mice. Scale bar: 50um. F. ELISA analysis of IL-1b protein in plasma of WT or Spp1KO-driven TPO fibrosis. n=3-5/group. G. Representative imaging of IL-1b staining and DAB quantification in 4um femoral sections TPO-driven WT or Spp1KO-driven fibrosis in WT recipient mice. Scale bar: 50um. H. mRNA expression of Spp1, Tnfa and Il-6 in WT or Spp1KO FL-HoxB8 cells (undifferentiated) carrying either the EV or MPLW515L mutation. n=3/group, P-value is based on a one-way ANOVA.(*) = p<0.05,(**)= p<0.01, (***) = p<0.001, (****) = p<0.0001. I. Flow cytometry analysis of macrophage differentiation (CD11b+ F4/80+ cells) from FL-HoxB8 cells (WT vs. Spp1KO, MPLW515L vs. EV) over 5 days, followed by PE+ latex bead phagocytosis assay performed for 1 hour. n=3/group, P-value is based on a one-way ANOVA.(*) = p<0.05, (****) = p<0.0001.

To better understand how the absence of Spp1 in hematopoietic progenitor cells and their monocyte/macrophage progeny affects inflammation and phagocyte function, we generated novel cells lines from WT C57BL/6J and Spp1KO mice: FMS-related tyrosine kinase 3 ligand (FL) estrogen-receptor (ER)-HoxB8 cells^24^, which are immortalized hematopoietic stem and progenitors cells with lymphoid and myeloid differentiation potential (FL-HoxB8). We transduced each cell line with either the MPLW515L-GFP or pMIG-EV-GFP retroviral vectors to obtain WT-EV, WT-MPL, Spp1KO-EV and Spp1KO-MPL FL-HoxB8 cells that can be maintained in a progenitor-like status with the addition of β-estradiol and Flt3 ligand. The MPLW515L mutation is common in patients with myelofibrosis and leads to activated JAK-STAT signaling. We confirmed in the WT MPL cells that STAT5b is overexpressed, as a surrogate marker for constitutive activation of the JAK-STAT pathway (Supplementary Fig. 5A). In both Spp1KO HoxB8 cell lines, regardless of mutational status, cells grew at a slower rate (Supplementary Fig. 5B). Interestingly, we demonstrated that the MPL mutation *per se* activates Spp1 expression which was not detectable in either Spp1KO cell lines, confirming the Spp1 knockout (Fig. 5H). In addition, the presence of the MPL mutation in HoxB8 cells, induces overexpression of pro-inflammatory markers *Tnfa* and *Il-6*, which were significantly downregulated in the Spp1KO, indicating Spp1 as a regulator of inflammation (Fig. 5H). To test the function of macrophages, we differentiated HoxB8 cells from the different lines into macrophages by withdrawing b-estradiol and supplementing media with m-CSF ligand. All lines readily differentiated into macrophages with comparable frequencies (Fig. 5I). However, when testing the function of macrophages evaluating the uptake of latex beads, the phagocytosis activity was increased in WT MPL mutant cells and restored to normal levels in the absence of Spp1 (Fig. 5I), indicating that Spp1 not only regulates the secretory phenotype of macrophages but also their phagocytic function.

### SPP1 is increased in the blood in MPN patients and high levels correlate with poor survival

To clearly understand the importance of SPP1 as a predictive marker of disease, we assessed levels of SPP1 protein in the plasma of MPN patients compared to healthy controls. We observed an increase in plasma SPP1 levels in MPN, irrespective of driver mutation (JAK2V617F vs. CALRdel52, Fig. 6A) compared to healthy controls. We next interrogated plasma proteomics and survival data from the UK Biobank, which contains de-identified biological samples and health-related data from participants aged 40-69, recruited between 2006 and 2010 across Great Britain. Given the chronic nature of myeloproliferative neoplasms (ICD10 code D47) and the potential latency between disease onset and formal diagnosis, we included participants who received a diagnosis within any time prior and within 2 years after the date of baseline assessment (i.e., blood draw). To reduce potential reverse causation and confounding by terminal illness, we excluded individuals who died within one year following the baseline visit and those without any SPP1-related data, leading to the inclusion of 33 samples with MPN (Supplementary Fig. 6A). We first compared the mean SPP1 concentrations (Normalized protein expression (NPX) units) between participants who died during follow-up (“died”) and those who remained alive (“survived”), and showed that higher SPP1 levels correlate death during follow-up (Fig. 6C). Critically, Kaplan–Meier curves demonstrated a stepwise decrease in survival with increasing SPP1 levels, with the highest mortality observed in Q4 (“high”, red) (Fig. 6D, Supp. Fig. 6B). Q1 (“low”, green) and Q2–Q3 (orange and yellow) show comparatively better survival over the 15-year follow-up period. We further investigated the diagnostic performance of SPP1 concentration in classifying patients with MPN who died versus those who survived during follow-up using Receiver Operating Characteristic (ROC) analysis and show that ROC curve rises in a stepwise manner, indicating that SPP1 concentration captures meaningful variation in mortality risk (Supplementary Fig. 6C). Altogether, these results indicate a clinical relevance of Spp1 as a prognostic marker in MPN patients.

**Figure 6.**
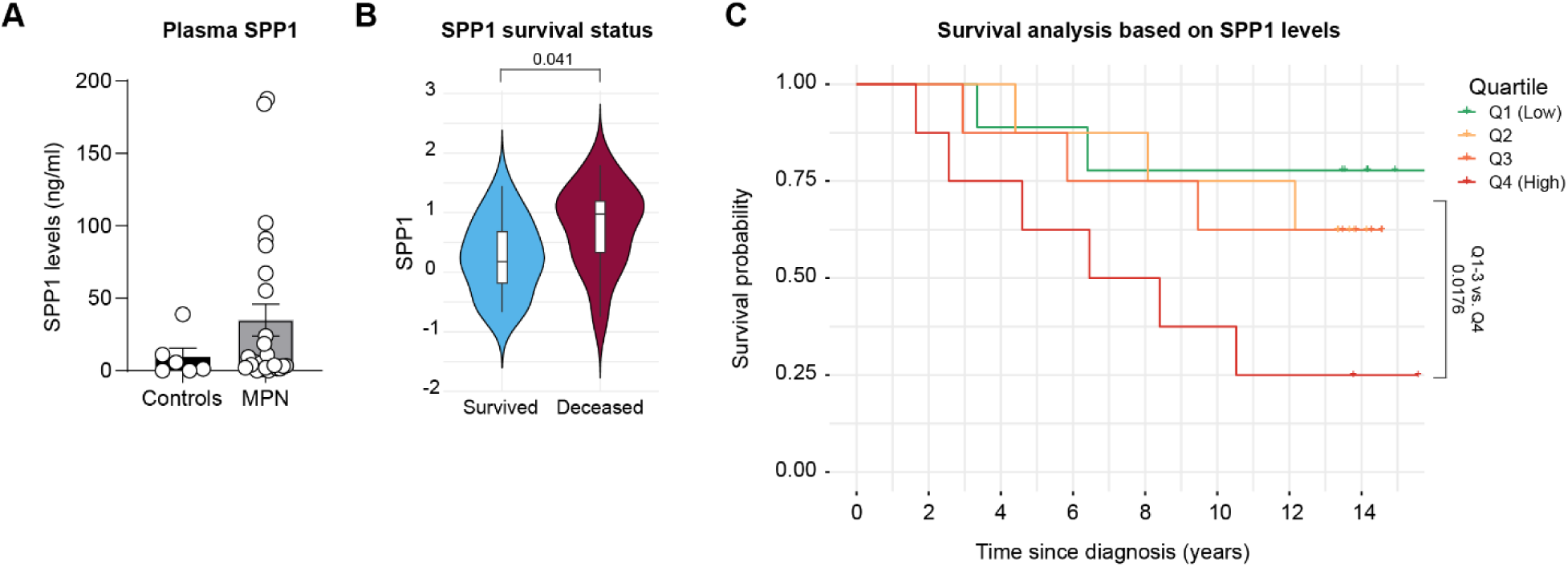
Elevated circulating SPP1 levels in MPN correlate with inferior survival outcomes. (A) ELISA analysis of SPP1 protein in human blood plasma of MPN patients and healthy controls. n=6 healthy, 15 MPN. Statistical test performed is an unpaired t-test, p=0.2796. (B) Normalized SPP1 plasma protein abundance in MPN patients for distinct outcomes. Differences between groups were evaluated using a student t-test. (C) Kaplan–Meier survival curves of primary myelofibrosis patients stratified by quartiles of SPP1 concentration at baseline assessment. Differences between groups were evaluated using the log-rank test.

## Discussion

Here, we identified SPP1-expressing macrophages as a pivotal component of the fibrotic BM microenvironment, functioning as a central “cellular glue” that links inflammatory resident-like and infiltrating macrophages with fibrosis-driving stromal cells through multimodal mechanisms of spatial proximity, extracellular matrix (ECM) communication, and cytokine-mediated activation.

Using an integrative approach combining dual lineage tracing, single-cell and multiplet sequencing, and our novel computational tool, NicheSphere, we demonstrate that Spp1+ macrophages are not merely passive bystanders but instead occupy a central communication hub within the fibrosis-interacting core niche. These macrophages exhibited preferential co-localization with osteoCAR cells, fibroblasts, megakaryocytes, and other hematopoietic subsets during disease progressions. Importantly, the functional importance of this macrophage subset was not identified with similar resolution and granularity in conventional pathway analyses for scRNA-seq analysis, underscoring the value of spatial co-localization-based analytics to uncover disease-relevant cellular interactions that may be missed by single cell transcriptional analysis alone.

NicheSphere uniquely identifies groups of cells that converge or diverge in disease by clustering them based on condition-specific differences in co-localization probabilities with specific partners. Compared to other state of the art tools, NicheSphere is the first approach allowing both the condition specific co-localization and communication analysis, which makes it unique. This approach optimizes the discovery of key interactions driving a given phenotype and enables multiple analytical strategies. NicheSphere prioritizes ligand–receptor (LR) pair-based interactions, providing an integrated database of ligands and associated biological processes, while also allowing users to supply custom curated LR sets (Supp. Table 1). This is in contrast to existing tools such as LIANA+^25^ and CellPhoneDB^26^, which are based on uncurated ligand-receptor (LR) databases and return around 18,000 significant LR and cell-cell interactions in either EV or TPO conditions. NicheSphere analysis, on the other hand, returns 9,785 significant interactions, which are further refined by pathways enriched in a particular condition and niche. For example, the TPO-enriched ECM glycoproteins and Collagens are related to 1,632 and 1,466 interactions respectively. Altogether, by combining co-localization and curation, NicheSphere improves interpretability by prioritizing and providing a mechanistic interpretation of predicted cell-cell interactions.

Among the spatially organized communication niches identified by NicheSphere, the “macrophage niche” (encompassing resident-like macrophages, Ly6C^low^ infiltrating macrophages, and dendritic cells) was characterized by enrichment in pro-inflammatory and cytokine-driven programs, including WNT, JAK-STAT, TNFα, and p53 signaling. In contrast, the “fibrosis-interacting core,” composed of osteoCAR cells, fibroblasts, megakaryocytes, and other stromal and hematopoietic subsets, exhibited upregulation of canonical pro-fibrotic pathways such as TGF-β and NF-κB, along with enhanced ECM glycoprotein signaling. Notably, Spp1+ macrophages were uniquely positioned at the intersection of these niches, exhibiting increased co-localization with both the macrophage-enriched compartments and the fibrosis-interacting core. This bridging role supports a model in which Spp1+ macrophages integrate inflammatory activation and stromal remodeling, effectively coordinating cytokine-driven inflammation with matrix deposition to drive disease progression.

Our results are consistent with findings in other fibrotic tissues, where Spp1+ macrophages have been implicated in ECM remodeling and immune regulation in the lung^19^, heart^21^ and kidney^20^. Our results indicate that SPP1 acts both as an adhesive ligand binding to integrins and CD44 on stromal cells, and as a signaling molecule shaping inflammatory phenotypes. Interestingly, SPP1^+^ macrophages were found to be closely associated with cancer associated fibroblasts (CAFs) in hypoxic tumor cores, and similarly reported to be actively communicating via integrin and CD44-mediated signaling, across cancer types^27,28^. Ligand-receptor mapping further revealed SPP1–integrin interactions as dominant communication axes, particularly involving osteoCAR and fibroblasts, which upregulated ECM glycoprotein and TGF-β signatures within the fibrosis-interacting core.

Functionally, genetic ablation experiments confirmed that stromal Spp1 is a driver of the fibrotic transformation. Loss of Spp1 in the stromal compartment ameliorated hallmark features of advanced fibrosis, anemia, splenomegaly, normalized bone marrow cellularity, and decreased reticulin deposition, highlighting that stromal-derived Spp1 is necessary to sustain the fibrotic niche. Complementary deletion of Spp1 in the hematopoietic compartment demonstrated that macrophage-derived SPP1 is likewise critical for inflammation and myeloproliferative features, as evidenced by the reduction of circulating IL-1β, monocytosis, and fibrosis severity. These bidirectional experiments underscore a reciprocal feed-forward loop wherein stromal and hematopoietic Spp1 cooperatively reinforce maladaptive remodeling.

Modeling of intracellular signaling and cell-cell communication revealed that Spp1+ macrophages propagate inflammatory activation via SPP1-mediated crosstalk with other macrophages, such as leading to IL-1β production and downstream stromal reprogramming. Notably, IL-1β induced both *Col1a1* and *Spp1* expression in stromal cells, while TGF-β promoted myofibroblast activation without upregulating *Spp1*, suggesting that IL-1β plays a more integrative role in inflammatory-fibrotic coupling, consistent with its known involvement in MPN^29^. The observation that Spp1 also regulates macrophage function is further interesting in the context of pathognomonic monocytosis in MPN and MF. Using HoxB8-derived macrophage progenitors, we showed that the MPL mutation induced both Spp1 expression and an inflammatory secretory phenotype marked by TNFα and IL-6 upregulation, while loss of Spp1 mitigated these effects and restored normal phagocytic activity. This implies that SPP1 not only orchestrates intercellular communication but also contributes to cell-autonomous macrophage activation and effector function in line with previous reports^30^, potentially contributing to the chronic inflammation and clearance of terminally differentiated myofibroblasts in the fibrotic core.

Together, our findings position Spp1+ macrophages as a central organizing element of the fibrotic bone marrow niche, acting through multimodal mechanisms: (1) promoting adhesive interactions with fibrosis-driving stromal cells, (2) stimulating pro-inflammatory cytokine production, and (3) modulating stromal differentiation programs. The consistent observation of Spp1/osteopontin as a central hub in the fibrosis-interacting core suggests that therapeutic targeting of the SPP1-integrin/CD44 axis could disrupt pathological cell-cell adhesion and niche activation. Our study additionally evaluates the prognostic potential of SPP1 in MPN, and we highlight here that plasma SPP1 levels are significantly elevated in MPN patients compared to healthy controls, regardless of driver mutations, suggesting a role in the MPN pathology. Higher baseline SPP1 concentrations were associated with increased mortality in MPN patients, suggesting that SPP1 could not only be used as a disease biomarker but also as a predictor of survival. Importantly, the absence of SPP1 ameliorated the MPN phenotype and fibrosis in mouse models, suggesting that it affects both the mutated hematopoietic clone and the fibrotic transformation. Our study suggests that (selectively) targeting SPP1—through biological or pharmacological approaches—may offer a promising pre-clinical strategy to address both the mutated clone and fibrosis in MPN patients.

Our data emphasize that focusing solely on cell-intrinsic transcriptional changes may underestimate the importance of spatially organized cell communities and ligand-receptor networks that shape disease phenotypes. Approaches integrating single-cell and multiplet sequencing, as implemented here with NicheSphere, represent a powerful framework to dissect these complex ecosystems. In summary, our study provides compelling evidence that Spp1+ macrophages serve as a communication hub integrating inflammatory and profibrotic programs in the bone marrow niche. By combining spatial and transcriptional analyses, we define SPP1 as a central mediator of pathological cross-talk and highlight a novel therapeutic axis in myeloproliferative neoplasm-associated fibrosis.

## Methods

### Mouse experiments

All mouse studies were approved by the Animal Welfare/Ethics committee of the EDC in accordance with legislation in the Netherlands (approval No. AVD1010020173387 and No. AVD101020216373). Gli1CreERt2 (i.e., Gli1tm3(re/ERT2)Alj/J, JAX Stock #007913), Rosa26tdTomato (i.e., B6-Cg-Gt(ROSA)26Sorttm(CAG-tdTomato)Hze/J JAX Stock #007909), Pf4-Cre mice (C57BL/6-Tg(Pf4-icre)Q3Rsko/J, JAX stock #008535) and the Spp1KO (B6.129S6(Cg)-Spp1tm1Blh/J, JAX stock #004936) mice were purchased from Jackson Laboratories (Bar Harbor, ME, USA). Offsprings were genotyped by PCR according to the protocol from the Jackson laboratory. CD45.1 mice were purchased from Envigo (B6.SJL-Ptprca Pepcb/BoyJ). Mice were maintained in specific-pathogen-free conditions, on a 12-hr light/dark cycle and were provided with water and standard mouse chow *ad libitum*. Mice were randomly assigned to experimental groups.

### Viral transductions

Lentiviral particles were produced by transient transfection with lentiviral plasmid together with pSPAX and VSVG packaging plasmids using TransIT (Mirus). Lentiviral particles were concentrated by ultracentrifugation at 4°C. For lentiviral transduction, CD117(c-Kit)-enriched cells from donor mice were isolated by crushing isolated pelvis, femurs, tibias, and subsequent CD117-enrichment by magnetic separation (Miltenyi Biotec). c-kit+ BM cells were cultured in StemSpan media (Stem Cell Technologies) supplemented by murine stem-cell factor (m-Scf, 50 ng/ml, Peprotech) and murine thrombopoietin (m-Tpo, 50 ng/ml, Peprotech). c-kit+ cells were transduced with virus (empty vector, EV-GFP or thrombopoietin-GFP (TPO-GFP) overexpression,) in the presence of polybrene (4 ug/ml) for 48 hours, then transplanted into recipient mice in sterile saline solution.

### Induction of myelofibrosis by overexpression of TPO in Pf4ZsGreen cells into Gli1tdTomato recipient mice

8-10-week-old Gli1CreERT2;tdTomato mice received four tamoxifen injections three weeks before irradiation and transplantation. Gli1-tdTomato recipients mice were lethally irradiated with 10.5Gy and received 4-5×10^5^ cells from 8-week-old Pf4-ZsGreen mice that had been harvested 48 hours prior to transplantation and transduced with TPO (TPO-BFP+) lentivirus or control empty vector lentivirus (EV-BFP+). Transplanted mice received drinking water supplemented with enrofloxacin (Baytril) for 3 weeks post-transplantation. Blood was periodically collected from recipient mice *via* submandibular bleeds into Microtainer tubes coated with K_2_EDTA (Becton Dickinson, NJ, USA) and complete blood counts were performed on a Horiba SciI Vet abc Plus hematology system. Mice were sacrificed when they had signs of advanced fibrosis as indicated by dropping hemoglobin levels and apparent weight loss at 56 days post-transplant.

At the end of the experiment, mice were sacrificed and BM cells were isolated from crushed pelvis and hind legs. Cells were labeled with directly conjugated antibodies, anti-mouse: Gr1, Ter119, CD3, B220, CD11b, ckit, CD41, F4/80, CD48, CD41, Sca1, CD45.2, and CD150 (all 1:100). All samples were analyzed by using an FACS LSRFortessa or FACS Aria III (BD Biosciences, San Jose, CA). Hoechst solution was added (1:10 000) to exclude dead cells, and data were analyzed by using FlowJo software version 10 (Tree Star Inc, Ashland, OR).

### Induction of myelofibrosis by overexpression of TPO in Spp1KO experiments

For *Spp1KO stromal experiments*, 4-5×10^5^ ckit+ (CD117+) cells isolated from CD45.1^+^ B6.SJL mice were transduced with TPO-GFP or its control vector EV-GFP, and were transplanted into lethally irradiated (10.5Gy) CD45.2^+^ C57BL/6J or Spp1KO recipients (n=4-5 mice/group). For *Spp1KO hematopoietic experiments*, 4-5×10^5^ ckit+ (CD117+) cells isolated from CD45.2^+^ C57BL/6J or Spp1KO mice were transduced with TPO-GFP or its control vector EV-GFP, and were transplanted into lethally irradiated (10.5Gy) CD45.1^+^ B6.SJL recipients (n=4-5 mice/group). Transplanted mice received drinking water supplemented with enrofloxacin (Baytril) for 3 weeks post-transplantation. Blood was periodically collected from mice *via* submandibular bleeds into Microtainer tubes coated with K_2_EDTA (Becton Dickinson, NJ, USA) and complete blood counts were performed on a Horiba SciI Vet abc Plus hematology system. Mice were randomly assigned to transplant groups and were sacrificed when mice showed signs of advanced fibrosis as indicated by dropping hemoglobin levels and apparent weight loss.

At the end of the experiment, mice were sacrificed and BM cells were isolated from crushed pelvis and hind legs. Cells were labeled with directly conjugated antibodies, anti-mouse: Gr1, Ter119, CD3, B220, CD11b, ckit, CD41, F4/80, CD48, CD41, Sca1, CD45.2, and CD150 (all 1:100). All samples were analyzed using an FACS LSRFortessa or FACS Aria III (BD Biosciences, San Jose, CA). Hoechst solution was added (1:10 000) to exclude dead cells, and data were analyzed by using FlowJo software version 10 (Tree Star Inc, Ashland, OR).

### Isolation of singlets and multiplets for scRNA-sequencing using 10X Genomics

After sacrifice, bones (femurs, tibia and hip bones) were crushed gently in PBS/2% FCS on ice (<5 minutes) and the cell solution was filtered through a 70μm nylon mesh using wide-bore pipette tips. BM cells were RBC lysed using PharmLyse buffer for 5 minutes, washed in PBS/2% FCS and kept on ice until further processing. The remaining bone chips were collected and digested for 15 minutes in 1mg/ml Collagenase II (Invitrogen #17101015) diluted in alphaMEM with 10% normal FCS and 1% P/S. After digestion, the bone chip fraction was washed in PBS/2% FCS and filtered through a 70μm nylon mesh followed by RBC lysis for 5 min. The bone chip fraction was then added to the previously isolated BM cells. Cells were resuspended in 300μl PBS/2% FCS and stained at 4°C for 15 min with the following antibodies: Gr1-APCCy7, Ter119-APCCy7, CD3-APCCy7, B220-APCCy7 and ZombieAqua for live/dead cells. Washing was performed by adding 1ml PBS/2% FCS and centrifuging for 5 min at 300×g, 4°C. After resuspension, TdTomato+ singlets, ZsGreen+ singlets or TdTomato+ZsGreen+ doublets from lineage/CD45/Ter119-negative cells were sorted into 50μl DMEM/10% FCS (BD Aria III) and used for the 10x platform. Unstained cells were used as negative controls to define gating. All antibodies were purchased from BioLegend.

### Single-cell library preparation and sequencing

For 10X sc-RNAseq processing, samples were loaded as follows: <u>sample 1</u> – TPO 10,000 ZsGreen^+^ droplets combined with 60,000 TdTomato^+^ droplets, <u>sample 2</u> – EV control 10,000 ZsGreen+ droplets combined with 60,000 TdTomato+ droplets, <u>sample 3</u> – TPO 60,000 double positive ZsGreen/TdTomato droplets, <u>sample 4</u> – EV control 60,000 double positive ZsGreen/TdTomato droplets. The libraries were prepared using the Chromium Single Cell 3′ Reagent Kits (v3.1): Single Cell 3′ Library & Gel Bead Kit v3.1 (PN-1000268), Chromium Next GEM Chip G Single Cell Kit v3.1 (PN-1000127) and Dual Index Kit TT Set A (PN-100215) (10x Genomics), and following the Chromium Single Cell 3’ Reagent Kits User Guide (v3.1 Chemistry Dual Index) (manual part no. CG000315, revision D). Finalized libraries were sequenced on a Novaseq6000 platform (Illumina), aiming for a minimum of 50,000 reads/cell using the 10x Genomics recommended number of cycles (28-8-0-91 cycles).

### Data QC and preprocessing

The scRNA-seq count matrices were obtained by aligning the raw sequencing reads into the custom td-tomato transcripts added mm10 mouse reference genome via the cellranger (version 4.0.0 and 6.0.2). Next, we used Seurat (v4.2) to analyze the scRNA-seq (Hao et al. 2021) based on R version 4.2.1. Cells with a high amount of mitochondrial genes (>20%), cells with low (<400) or high feature counts (>40,000) were filtered out, and the Xist gene was removed to mitigate gender differences in downstream analyses. Cell-cycle related genes, the proportion of mitochondrial, ribosomal, and UMI counts were regressed out and a log-normalization of read counts was performed. Singlets and multiplet scRNA-seq libraries were separately processed as follows: Condition specific scRNA-seq libraries were integrated with Harmony^31^ using the first 50 components from a PCA computed using Seurat^32^. Shared nearest neighbor (SNN) based clustering was performed with Seurat using the first 50 components of the dimensionality reduction obtained through Harmony integration. Cell-cell communication analysis was performed on the clustered results using the CellphoneDB method implemented in Liana^18^ with default parameters. Intracellular signaling analysis was performed with scSeqCommDiff framework^33^ with default parameters, comparing TPO vs EV conditions across the respective clusters.

### Estimating multiplets cell composition using deconvolution

Celltype deconvolution was performed on the multiplets using the R library BayesPrism^14^, which models prior distribution using scRNA-seq, and leverages each observed bulk data to infer a joint posterior distribution of cell proportion and gene expression in bulk samples. BayesPrism was employed to predict cell type proportions in multiplets. Before cell type deconvolution, gene filtering was performed using the function ‘cleanup.genes’. Ribosomal and mitochondrial genes were removed. Genes on chromosome X and Y were also removed to prevent sex-specific differences between the singlets and the multiplets. Lowly expressed genes were removed by selecting genes expressed in at least 5 cells, as performed in Chu et al., 2022^14^. Signature genes were selected by performing pairwise t-test between different cell types using the function ‘get.exp.stat’. The test was performed on cell types of which more than 20 cells were available. To run the deconvolution step, the singlets data was subsetted, taking the genes with a p-value<0.01 and a minimum log fold change of 0.1 using the ‘select.marker’ function. We used the function ‘new.prism’ to combine the singlets and multiplets data together into a prism object. We used our subset singlets dataset as input for this function, as well as a subset version of the multiplets dataset with the same genes. Then we filtered genes whose expression fraction was greater than 0.01 in more than 10% of the multiplets (outlier.cut=0.01, outlier.fraction=0.1). Finally, we ran the BayesPrism model using the function ‘run.prism’ and extracted the posterior mean of cell type fractions using the function ‘get.fraction’.

### NicheSphere

Nichesphere is a framework for cellular interaction aware cell-cell communication inference. It consists of the following main steps: 1) Estimation of the co-localization probabilities cell types; 2) Condition specific cell-cell colocalization enrichment and networks; 3) Co-localization and process constrained cell-cell communication. Nichesphere requires as input a clustered annotated singlet cell RNA, a cell deconvolution results by mapping the singlet to the multiplet data using any cell deconvolution algorithm and a singlet based ligand-receptor cellular communication analysis (see above for details of the cell deconvolution and LR analysis tools used in this study). Cell types and conditions (disease vs normal) of the deconvolution and cell-cell communication needs to match. Another novel feature of NicheSphere is the curation of ligand-receptor pairs in cell-cell communication in ten distinct mechanisms related to extracellular matrix (ECM)^34^ and immune recruitment signaling^35^ (Supp. Table 1). Moreover, these LR sets are mostly related to exclusive sets of LR pairs (Supp. Fig. 2F). A formal description of NicheSphere is provided in the Supplementary File 1.

Of note, previous computational workflows for PIC-seq data, for example, were based on the use of customized scripts for the analysis of a single condition data set^10^. Their approach was simply based on contrasting expected cell type numbers pairs by considering singlet cell proportions with the number of doublet cells detected in the PIC-seq experiment of the corresponding sample. Later, CIM-seq improved the previously described procedure by including statistical approaches to find condition specific enrichment interaction^11^. Both approaches are not able to contrast co-localization interactions between two conditions; or to relate cell-cell communication mechanisms associated with these. NicheSphere is to the best of our knowledge the only approach to infer condition specific niches from co-localization and communication networks. Moreover, it is the only method implemented as a tool and python package available in Github (https://github.com/CostaLab/Nichesphere) supporting its use by the community.

### Histological and immunohistological analysis

Murine organs were fixed in 4% paraformaldehyde for 24 hours and transferred to 70% ethanol. Spleens were weighed before fixation. Femurs were decalcified in 10% EDTA/Tris-HCl (pH 6.6) solution for 6 days, dehydrated, and paraffin embedded according to standard pathology protocols. H&E staining was performed according to established routine protocols and reticulin staining was performed using the Reticulin silver plating kit according to Gordon & Sweets (Merck, #1.00251) on 4 μm sections. For IL-1b and Spp1 staining, antigen retrieval was performed using citrate-buffer in a conventional lab microwave (Vector, antigen unmasking solution). Sections were treated with 3% H2O2 and blocked with Avidin/Biotin blocking kit (Vector), and subsequently incubated with primary antibody (goat anti-Spp1: AF808, Abcam, 1:200; rabbit-anti-Il-1b: P420B, Thermo Fisher, 1:150) overnight at 4°C. Biotinylated anti-rabbit-antibody or anti-goat-antibody (Vector) was used as a secondary antibody for 30 min at room temperature. Slides were incubated with AB complex for 30 min at room temperature, washed, and incubated for a further 10 min with DAB substrate. Slides were counterstained with hematoxylin and mounted with glass coverslip using DPX mounting media (Sigma). Slides were scanned and digitized in an automated fashion using a Hamamatsu Nanozoomer 2.0 HT system. Images were analyzed and exported using the NDP.view software (Hamamatsu, V2.5.19).

### Sample processing for immunofluorescence imaging

At the time of sacrifice, murine long bones (femur and/or tibia) were harvested and fixed immediately in ice-cold 2% paraformaldehyde (PFA) for 6–8 h under gentle agitation. Bones were decalcified in 0.5 M EDTA for 16–24 h at 4°C under gentle shaking agitation, followed by overnight incubation in cryopreservation solution (20% sucrose, 2% PVP) and embedding in bone embedding medium (8% gelatine, 20% sucrose, 2% PVP), as previously described^36,37^. Samples were stored overnight at −80 °C. 60μm-thick cryosections were prepared for immunofluorescence staining. Bone sections were washed in ice-cold PBS and permeabilized with ice-cold 0.3% Triton-X-100 in PBS for 10 mins at room temperature (RT). Samples were incubated in blocking solution (5% heat-inactivated donkey serum and 0.1% Tween20 in PBS) for 30 min at RT. Primary antibodies (Anti-CD68 antibody [EPR23917-164, Abcam ab283654, 1:100], anti-TREM2 antibody [Abcam, ab95470, 1:100], anti-CD9-FITC [Biolegend, 124807, 1:100] and anti-Lipoprotein Lipase/LPL Antibody-AF700 [Novus Biologicals, NBP2-71178AF700, 1:100]) were diluted in 5% donkey serum, 0.1% Tween20 in PBS and incubated overnight at 4°C with DAPI (1:1,000). Slides were washed 3–5 times in PBS in 5–10 min intervals. Species-specific Alexa Fluor secondary antibodies Alexa Fluor 488 (Thermo Fisher Scientific, Cat#A21208), Alexa Fluor 546 (Thermo Fisher Scientific, Cat#A11056), Alexa Fluor 594 (Thermo Fisher Scientific, Cat#A21209), Alexa Fluor 647 (Thermo Fisher Scientific, Cat#A31573 or Cat#A21447) diluted 1:400 in PBS were added and incubated for 2 h at RT. Samples were embedded in Fluoromount-6 (Cat#F3724, ITK Diagnostics) and imaged within 3 days. Confocal microscopy was performed with a Leica SP8 DLS equipped with the following lasers: 405nm, argon 488nm, 561nm and 633nm and a HC PL APO CS2 objective (Leica).

### SPP1 ELISA in human plasma samples

MPN patient plasma samples were collected and supplied by the University Clinic Hamburg-Eppendorf (UKE), obtained via the German Study Group MPN Aachen and the Department for Molecular Diagnostics at the Erasmus Medical Center (MC) in Rotterdam. Control samples were obtained via the Institute for Clinical Chemistry at Erasmus MC. Surplus material was collected from dermatological, and cardiological, orthopedic or neurological patients after diagnostics according to the ethical vote MEC-2018-1445 and processed on the day of acquisition. All subjects had no history or indication of any hematological or non-hematological malignancy. Samples were de-identified at the time of inclusion. All patients provided informed consent and the data collection was performed in accordance with the Declaration of Helsinki. Plasma was isolated from whole blood anticoagulated with EDTA by centrifugation (2000 × g, 7 min) and stored at −80°C. Samples were thawed on ice and centrifuged for 5 min at 2500 × g before further processing. Samples were diluted 1:50 and SPP1 concentration was quantified using the Human SPP1 Quantikine ELISA (R&D Systems, Human Osteopontin DOST00) according to the manufacturer’s instructions.

### Generation and cell culture of WT-EV, WT-MPL^W515L^, Spp1KO-EV and Spp1KO-MPL^W515L^ Hoxb8-FL progenitors cell lines

Bone marrow cell isolation was performed as described previously. Isolated bone marrow cells from C57BL/6 or Spp1KO mice were resuspended at a concentration of 5×10^5^ cells/ml in IMDM medium (GIBCO) with recombinant mouse IL-3 (10 ng/ml), IL-6 (20 ng/ml), and 50 ng/ml recombinant SCF (R&D system). After 2 days of cell culture, cells were seeded at a concentration of 2×10^5^/ml per well in a 12-well plate using progenitor outgrowth medium (POM), which consists of Iscove’s modified Dulbecco’s medium (IMDM) supplemented with 10% FBS, 1% penicillin-streptomycin (GIBCO), 1 µM β-estradiol (Sigma), and 25 ng/ml recombinant mouse Flt-3 ligand for generation of Hoxb8-FL cells^24^. Cells were infected with MSCV-ERHBD-Hoxb8 vector by spin inoculation. The MSCV-ERHBD-Hoxb8 plasmid was a generous gift from Hans Haecker (St. Jude Faculty, Department of Infectious Diseases, St. Jude Children’s Research Hospital, Memphis, USA). Once established, Hoxb8-FL cell lines were transduced with pMIG-MPL^W515L^ or pMIG-EV retroviral vectors and sorted on GFP+ expression to establish pure cultures. HoxB8-FL cells were maintained in IMDM medium (GIBCO) supplemented with 10% FBS, 1% P/S, 40ng/ml recombinant mouse Flt-3 ligand (Peprotech), and 1 µM β-estradiol (Sigma).

### RNA extraction and real-time qPCR analysis

RNA extraction and real-time quantitative PCR analysis RNA from pelleted MSCs or FL-HoxB8 cells were extracted using Trizol solution (Thermo Fisher) according to the manufacturer’s instructions, and 1μg of total RNA was reverse transcribed with Superscript IV (Invitrogen). Total RNA was reverse transcribed using the high-capacity cDNA Reverse Transcription kit (Applied Biosystems). Quantitative polymerase chain reactions were performed with SYBRGreen PCR master mix (Thermo Fisher) on an Applied Biosystems 7500 Real-Time PCR System. Glyceraldehyde-3-phosphate dehydrogenase (*Gapdh*) was used as a housekeeping gene. Data were analyzed using the 2-ΔΔct method.

### Study population and case definition

We used data from the UK Biobank, a prospective cohort study comprising approximately 500,000 participants aged 40–69 years at baseline (2006–2010), with linkage to national health records and biobanked blood samples. For this analysis, we identified participants with a diagnosis of myelofibrosis based on the first occurrence of ICD-10 code D47 (“neoplasms of uncertain or unknown behavior of lymphoid, hematopoietic and related tissue”) in hospital inpatient or death registry data.

Given the chronic nature of myelofibrosis and the potential latency between disease onset and formal diagnosis, we included participants who received a D47 diagnosis any time prior and up to 2 years after the date of baseline assessment (i.e., blood draw), and for which SPP1 protein data was available. To reduce potential reverse causation and confounding by terminal illness, we excluded individuals who died within one year following the baseline visit. Finally, 33 patients with a diagnosis of D47 and available measurements of SPP1 were included in the analyses.

### Protein biomarker measurement

Circulating levels of SPP1 (Secreted Phosphoprotein 1, also known as osteopontin) were quantified in EDTA plasma samples collected at baseline, using Olink® proximity extension assays (cardiometabolic and inflammatory panels). Protein expression values are reported on a log2-transformed NPX (Normalized Protein eXpression) scale, which allows relative quantification across participants. Participants were stratified into quartiles (Q1–Q4) based on SPP1 levels in the overall UK Biobank proteomics subsample, with Q1 representing the lowest and Q4 the highest protein concentrations.

### Survival analysis

Time-to-event analysis was performed to estimate all-cause mortality following baseline in D47-diagnosed individuals. The index date was defined as the baseline assessment date, where the blood was taken. Survival time was calculated in years from baseline until death or end of follow-up (censoring date: 1^st^ of January 2024). Kaplan–Meier survival curves were generated for each SPP1 quartile group. Differences between groups were evaluated using the log-rank test. All statistical analyses were conducted using R (version RStudio 2022.07.1), and visualizations were produced using the survival and survminer packages.

### Comparative analysis across SPP1 quartiles

To contextualize survival differences and explore potential correlates of SPP1 levels, we conducted descriptive and comparative analyses of demographic, clinical, and biochemical variables across SPP1 quartiles. Variables included age, sex, body mass index (BMI), renal and hepatic function markers, lipid parameters, glucose metabolism indicators, inflammatory markers, and circulating proteins. Continuous variables are presented as mean ± standard deviation, and comparisons across quartiles (Q1–Q4) were assessed using one-way analysis of variance (ANOVA). We additionally conducted pairwise comparisons between Q1 and Q4, as well as between Q1–Q3 vs Q4, to examine potential threshold effects of high SPP1 levels. P-values were reported for overall group differences and key pairwise contrasts, with nominal significance defined as *p* < 0.05.

### SPP1 group comparison by mortality status

To investigate the association between baseline SPP1 levels and vital status, we stratified participants by all-cause mortality (death vs. survival) based on linkage to the UK Biobank death registry. Baseline SPP1 levels were measured using Olink® proximity extension assays and are expressed in NPX (Normalized Protein Expression) units on a log2 scale. We calculated mean SPP1 levels and corresponding 95% confidence intervals separately for individuals who died and those who survived. Group differences were evaluated using an unpaired two-sample t-test, assuming unequal variances. The resulting p-value was considered nominally significant at the 0.05 level. All analyses and visualizations were performed using R (version 2022.07.1) and the ggplot2 package.

### Receiver operating characteristic (ROC) analysis for prediction of all-cause mortality

To evaluate the predictive performance of the selected biomarker for all-cause mortality, we computed a receiver operating characteristic (ROC) curve. The binary outcome variable was defined as death during follow-up (yes/no), based on linkage to national death registry data in the UK Biobank. The predictor variable was SPP1 assessed at baseline. We used the pROC package in R (version 2022.07.1) to compute the ROC curve and calculate the corresponding area under the curve (AUC). The analysis was performed on complete cases only. Sensitivity and specificity were derived across a range of thresholds, and the stepwise nature of the ROC reflects the number of events (deaths) available in the dataset.

### Statistical analysis

Statistical analysis was performed by using GraphPad Prism version 10 software (GraphPad Software Inc, San Diego, CA). Comparisons between 2 groups were performed by using an unpaired Student t-test or Mann-Whitney U test as described in the figure legends. For multiple group comparisons, an analysis of variance with post-hoc Tukey correction or a Kruskal-Wallis test was applied. Data are shown as mean ± standard error of the mean, and a value of P <.05 was considered significant.

## Supplemental tables and figures

**Supplementary table 1.** Biological processes database.

| Process | Database | Number of ligands | Number of associated LR pairs (from lianaDB consensus) |
| --- | --- | --- | --- |
| Collagens | Matrisome (PMID:36399478) | 45 | 321 |
| ECM affiliated | Matrisome (PMID:36399478) | 129 | 143 |
| ECM glycoproteins | Matrisome (PMID:36399478) | 168 | 422 |
| ECM regulators | Matrisome (PMID:36399478) | 204 | 195 |
| Proteoglycans | Matrisome (PMID:36399478) | 36 | 43 |
| SecretedFactors | Matrisome (PMID:36399478) | 265 | 1225 |
| Chemokine | Cytosig (PMID:34594031) | 42 | 280 |
| Cytokine | Cytosig (PMID:34594031) | 86 | 237 |
| GrowthFactor | Cytosig (PMID:34594031) | 130 | 1064 |
| Inhibitory | Cytosig (PMID:34594031) | 36 | 49 |

**Supplementary table 2.** Analysis of SPP1 quartiles.

| Variable | Q1.Q1 | Q2.Q2 | Q3.Q3 | Q4.Q4 | p_over all | p_Q1_vs_Q4 | p_Q13_vs_Q4 |
| --- | --- | --- | --- | --- | --- | --- | --- |
| Survival [years] | 12.23<br>± 4.30 | 11.61<br>± 3.51 | 11.12<br>± 4.53 | 7.94 ±<br>5.08 | 0.222 | 0.0837 | 0.087 |
| Time to death [days] | 4828.5<br>6 ±<br>2268.4<br>2 | 4400.0<br>0 ±<br>1516.4<br>4 | 4483.2<br>5 ±<br>1388.1<br>7 | 4368.0<br>0 ±<br>2243.4<br>8 | 0.956 | 0.68 | 0.811 |
| Event death [0/1] | 0.22 ±<br>0.44 | 0.38 ±<br>0.52 | 0.38 ±<br>0.52 | 0.75 ±<br>0.46 | 0.172 | 0.0303 | 0.0421 |
| BMI [kg/m<sup>2</sup>] | 25.83<br>± 4.92 | 27.39<br>± 4.88 | 31.27<br>± 5.27 | 30.33<br>± 8.30 | 0.233 | 0.208 | 0.489 |
| Age [years] | 55.78<br>± 8.83 | 63.50<br>± 3.96 | 57.38<br>± 6.37 | 58.00<br>± 9.65 | 0.208 | 0.629 | 0.842 |
| Sex [% female] | 0.22 ±<br>0.44 | 0.75 ±<br>0.46 | 0.62 ±<br>0.52 | 0.62 ±<br>0.52 | 0.143 | 0.108 | 0.626 |
| Alanine amino transferase [U/L] | 25.47<br>±<br>14.96 | 23.62<br>±<br>18.20 | 27.09<br>± 7.73 | 39.98<br>±<br>37.07 | 0.447 | 0.328 | 0.31 |
| Albumin [g/L] | 44.48<br>± 2.16 | 42.22<br>± 3.23 | 42.02<br>± 4.31 | 44.05<br>± 4.38 | 0.429 | 0.808 | 0.515 |
| Alkaline phosphatase [U/L] | 75.40<br>±<br>37.25 | 95.70<br>±<br>31.71 | 105.46<br>±<br>44.43 | 127.06<br>±<br>58.46 | 0.133 | 0.054 | 0.142 |
| ApolipoproteinA [g/L] | 1.32 ±<br>0.20 | 1.34 ±<br>0.19 | 1.45 ±<br>0.21 | 1.25 ±<br>0.18 | 0.251 | 0.487 | 0.14 |
| ApolipoproteinB [g/L] | 0.95 ±<br>0.20 | 0.98 ±<br>0.44 | 0.98 ±<br>0.24 | 1.04 ±<br>0.23 | 0.927 | 0.393 | 0.46 |
| Aspartate amino transferase [U/L] | 33.26<br>±<br>11.08 | 28.70<br>±<br>13.32 | 28.45<br>± 6.29 | 40.83<br>±<br>16.14 | 0.191 | 0.313 | 0.143 |
| C reactive protein [mg/L] | 2.74 ±<br>2.07 | 3.51 ±<br>3.35 | 4.33 ±<br>2.54 | 11.61<br>±<br>18.57 | 0.225 | 0.22 | 0.257 |
| Calcium [mmol/L] | 2.32 ±<br>0.11 | 2.37 ±<br>0.10 | 2.34 ±<br>0.14 | 2.32 ±<br>0.10 | 0.731 | 0.985 | 0.523 |
| Cholesterol [mmol/L] | 5.12 ±<br>1.17 | 5.51 ±<br>1.92 | 5.13 ±<br>1.00 | 5.50 ±<br>1.05 | 0.881 | 0.492 | 0.59 |
| Creatinine [μmol/L] | 63.72<br>±<br>14.39 | 85.15<br>±<br>24.93 | 74.69<br>±<br>17.66 | 100.11<br>±<br>40.61 | 0.0484 | 0.041 | 0.119 |
| CystatinC [mg/L] | 1.01 ±<br>0.29 | 1.08 ±<br>0.20 | 1.01 ±<br>0.26 | 1.41 ±<br>0.41 | 0.036 | 0.0422 | 0.0387 |
| Direct bilirubin [μmol/L] | 1.89 ±<br>0.67 | 2.07 ±<br>0.64 | 2.10 ±<br>0.49 | 2.50 ±<br>1.61 | 0.711 | 0.514 | 0.598 |
| Gamma glutamyl transferase | 32.98<br>±<br>29.17 | 54.59<br>±<br>53.84 | 48.30<br>±<br>32.55 | 114.11<br>±<br>87.68 | 0.0303 | 0.0359 | 0.0624 |
| Glucose [mmol/L] | 4.76 ±<br>0.78 | 5.29 ±<br>1.34 | 6.23 ±<br>3.11 | 4.76 ±<br>0.47 | 0.313 | 0.998 | 0.145 |
| HbA1c [mmol/mol] | 33.09<br>± 4.03 | 34.65<br>± 5.69 | 41.96<br>±<br>21.06 | 32.80<br>± 4.84 | 0.442 | 0.922 | 0.312 |
| HDL cholesterol [mmol/L] | 1.26 ±<br>0.27 | 1.28 ±<br>0.23 | 1.34 ±<br>0.33 | 1.11 ±<br>0.21 | 0.353 | 0.22 | 0.0602 |
| IGF.1 [nmol/L] | 21.18<br>± 6.87 | 24.91<br>±<br>15.93 | 15.22<br>± 5.92 | 18.67<br>± 6.38 | 0.249 | 0.447 | 0.571 |
| LDL direct [mmol/L] | 3.22 ±<br>0.84 | 3.57 ±<br>1.45 | 3.23 ±<br>0.77 | 3.56 ±<br>0.77 | 0.81 | 0.4 | 0.524 |
| Lipoprotein A [nmol/L] | 48.39<br>±<br>47.58 | 55.94<br>±<br>80.46 | 37.94<br>±<br>31.60 | 22.65<br>±<br>18.99 | 0.656 | 0.172 | 0.0929 |
| Phosphate [mmol/L] | 1.26 ±<br>0.18 | 1.23 ±<br>0.16 | 1.07 ±<br>0.17 | 1.16 ±<br>0.18 | 0.163 | 0.337 | 0.796 |
| SHBG [nmol/L] | 81.59<br>±<br>62.47 | 46.96<br>±<br>16.32 | 57.26<br>±<br>28.95 | 63.15<br>±<br>21.44 | 0.315 | 0.451 | 0.916 |
| Testosterone [nmol/L] | 5.07 ±<br>7.16 | 8.69 ±<br>5.22 | 6.13 ±<br>4.20 | 10.74<br>± 6.04 | 0.28 | 0.15 | 0.179 |
| Total bilirubin [μmol/L] | 8.70 ±<br>3.66 | 9.24 ±<br>3.35 | 11.27<br>± 5.68 | 8.69 ±<br>5.45 | 0.641 | 0.995 | 0.663 |
| Total protein [g/L] | 75.12<br>± 4.41 | 72.76<br>± 4.05 | 77.75<br>± 7.04 | 75.21<br>± 7.97 | 0.458 | 0.978 | 1.0 |
| Triglycerides [mmol/L] | 1.38 ±<br>0.58 | 1.41 ±<br>0.60 | 1.48 ±<br>0.81 | 2.34 ±<br>1.23 | 0.0878 | 0.0742 | 0.0784 |
| Urate [μmol/L] | 247.56<br>±<br>48.10 | 354.71<br>±<br>54.46 | 279.18<br>±<br>106.43 | 444.45<br>±<br>204.30 | 0.0121 | 0.0298 | 0.071 |
| Urea [mmol/L] | 5.08 ±<br>1.29 | 6.80 ±<br>2.55 | 4.43 ±<br>0.60 | 7.30 ±<br>2.02 | 0.00714 | 0.0209 | 0.0397 |
| Vitamin D | 58.27<br>±<br>26.22 | 43.24<br>±<br>16.86 | 50.63<br>±<br>22.40 | 42.40<br>±<br>37.65 | 0.638 | 0.346 | 0.555 |
| Townsend deprivation index at recruitment [index] | -0.60 ±<br>2.27 | -1.62 ±<br>2.86 | -1.20 ±<br>3.58 | -0.56 ±<br>3.11 | 0.867 | 0.976 | 0.661 |

**Supplementary figure 1.**
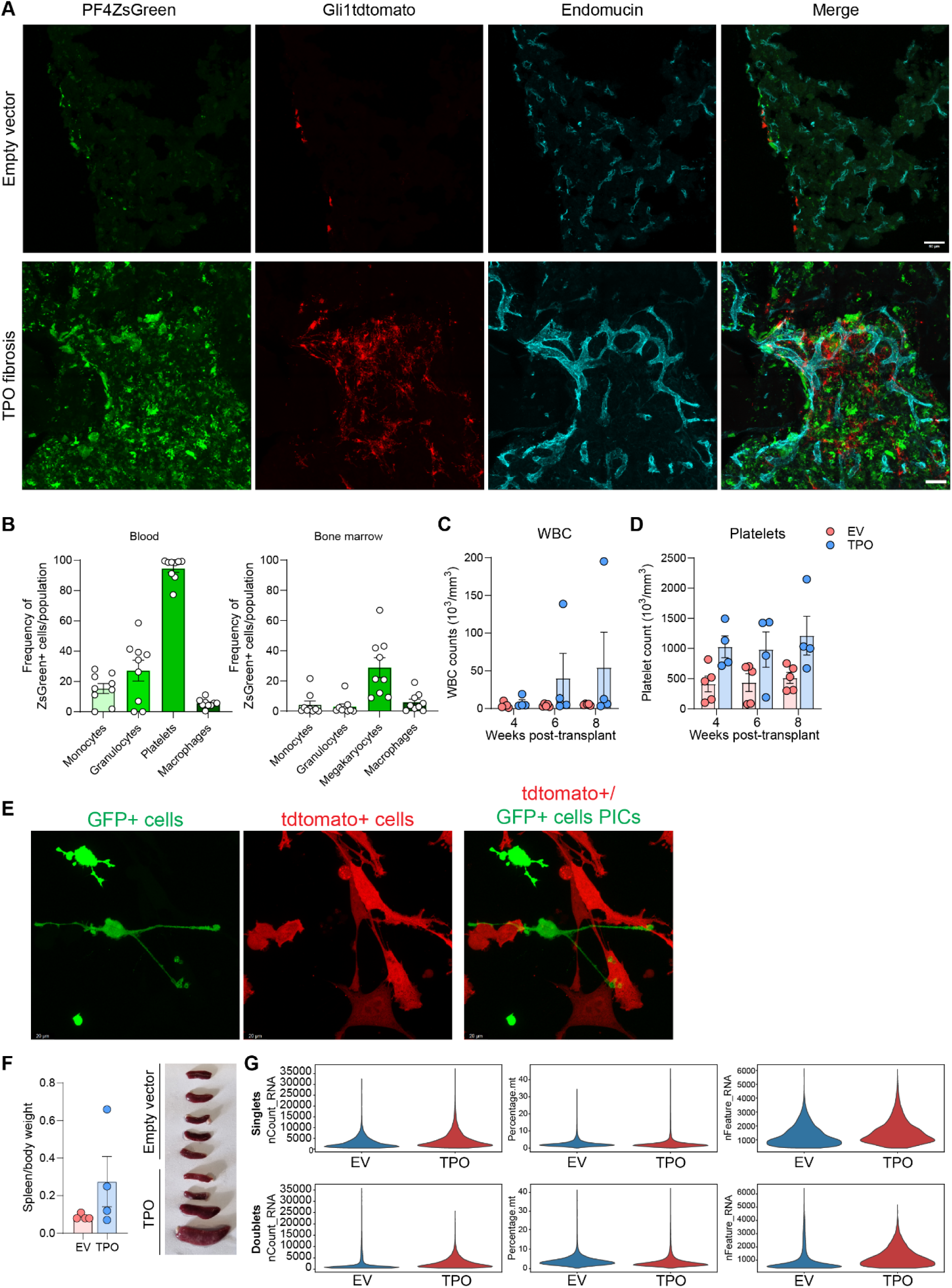
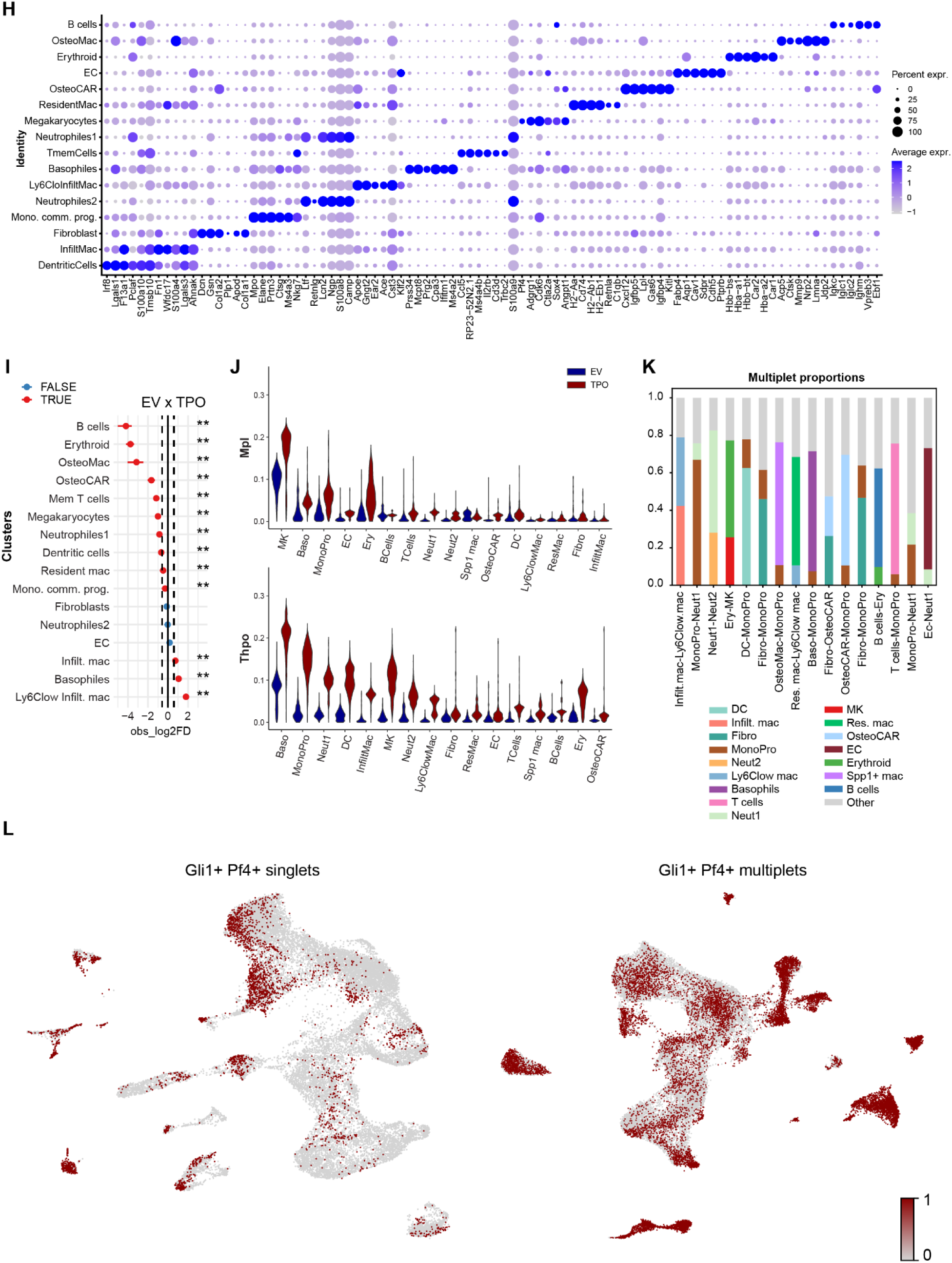
Mapping TPO-Induced cell–cell interactions in primary myelofibrosis using PICseq. (A) Confocal imaging of Pf4-ZsGreen (green) cells transplanted into Gli1-tdTomato+ (red) recipient mice, overexpressing either TPO or the empty vector (BFP), co-stained with endomucin for vasculature (cyan). Scale bar: 50um. (B) Myeloid lineage analysis of ZsGreen+ cells in blood and bone marrow (BM) using flow cytometry in mice transplanted with ZsGreen+ cells (TPO– and EV-transduced). Cell populations are gated as follows: monocytes: live CD45+CD11b+Gr1-F4/80-, granulocytes: live CD45+CD11b+Gr1+, platelets/megakaryocytes: live CD45+CD11b-Gr1-F4/80-CD41+, macrophages: live CD45+CD11b+Gr1-F4/80+. (C) White blood cell counts in TPO-driven fibrosis in Gli-TdTomato recipient mice with PF4-ZsGreen BM at the time of sacrifice. (D) Platelet counts in TPO-driven fibrosis in Gli-TdTomato recipient mice with PF4-ZsGreen BM at the time of sacrifice. (E) Representative images of physically interacting bone-marrow derived PF4-Zs (GFP) and Gli1tdtomato+ stromal cells. Scale bar: 20um. (F) Spleen-to-bodyweight ratio and spleen images in TPO-driven fibrosis in Gli-TdTomato recipient mice with PF4-ZsGreen BM at the time of sacrifice. (G) Sequencing data QC. Violin plots showing quality measures distributions across singlets (top panels) and multiplets (bottom panels) of each condition: Left panels show number of transcript counts; middle panels show percentage of mitochondrial counts; right panels show number of features detected (genes) (H) Dot plot showing expression of known BM cell population marker genes in the scRNA-seq singlet clusters (I) Log_2_(TPO/EV) fold-change in the number of cells of each type. Confidence intervals (95%) for the magnitude difference are based on bootstraping (n=1000) with scproportiontest. (J) Violin plots showing expression of Mpl receptor and Thpo ligand in TPO (red) and EV (blue) singlet populations. (K) Multiplet cluster compositions, based on cell signatures identified in the singlet populations with Bayesprism^14^. (L) UMAPs displaying Pf4+Gli1+ singlets and Pf4+Gli1+ multiplets, previously sorted for ZsGreen and tdTomato from TPO and EV control mice and sequenced using the 10X Genomics pipeline, n=27,046 singlets, n=35,927 multiplets.

**Supplementary figure 2.**
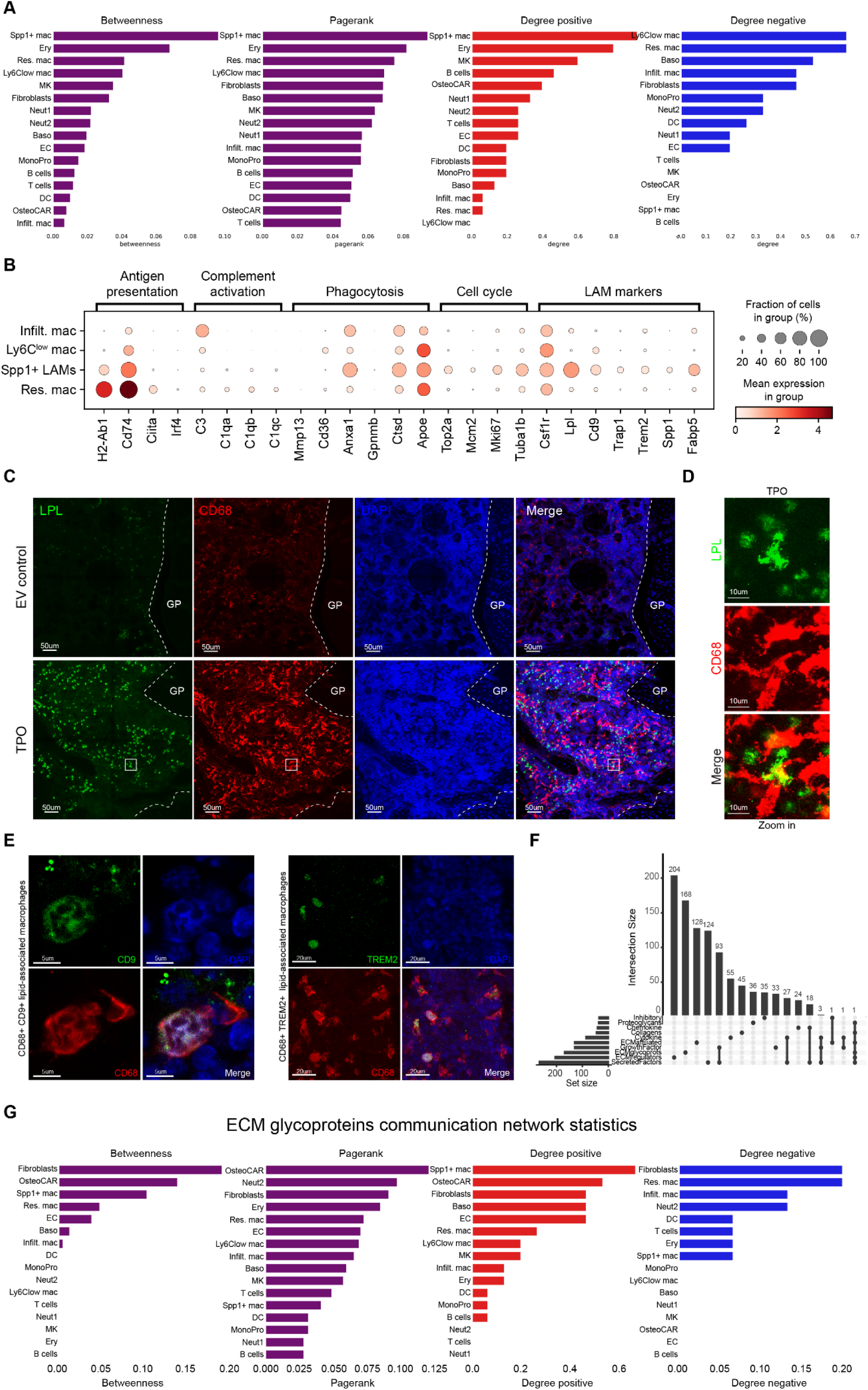
Multimodal characterization of macrophage subsets and fibrosis-associated ligand networks identified using NicheSphere in TPO-induced BM fibrosis. (A) Co-localisation network statistics, including betweenness, pagerank and positive/negative degrees. (B) Dot plot showing normalised gene expression of lipid-associated macrophages (LAMs) and specific macrophage modules^17^, in the scRNA-seq singlet dataset. Dot size represents the percentage of cells expressing the gene, and color represents the scaled mean gene expression of the cluster. (C) Representative imaging of CD68 and Lpl staining in 60um femoral sections of EV– or TPO-induced BM. Scale bar: 50um, 40X. White square indicates the zoom-in area used for panel D. (D) Zoom-in images of TPO-induced BM with co-localised staining of CD68 and LPL. Scale bar: 10um. (E) Left panels: Representative imaging of CD68+TREM2+ macrophages, in 40um femoral sections of TPO-induced BM. Scale bar: 20um, 40X. Right panels: representative imaging of CD68+CD9+ macrophages, in 40um femoral sections of TPO-induced BM. Scale bar: 5um, 63X. (F) Upset plot showing size and overlap among ligand sets in the fibrosis specific NicheSphere database listed in Supplementary Table 1. This indicates that most categories have non-overlapping LR pairs. (G) Extracellular matrix glycoproteins communication network statistics, including betweenness, pagerank and positive/negative degrees.

**Supplementary figure 3.**
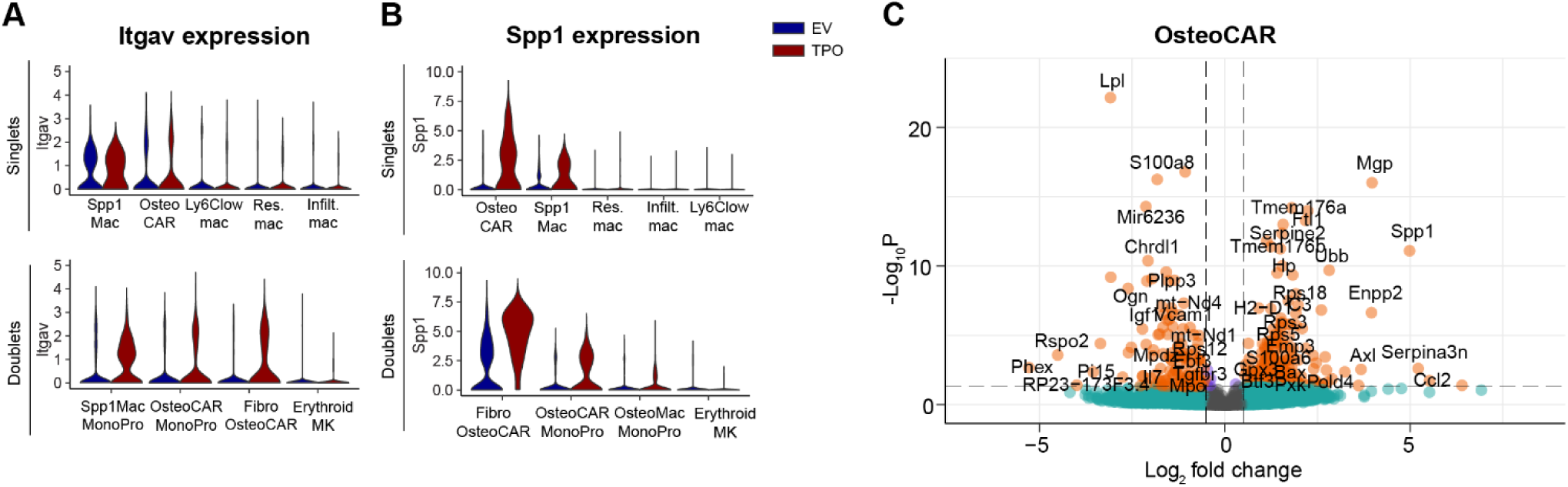
Differential expression analysis points towards fibrosis specific cell-cell interactions through Spp1 and Itgav in the stromal compartment. (A) Violin plots showing expression of Itgav in TPO (red) and EV (blue) singlets (upper panel) and doublets (lower panel). (B) Violin plots showing expression of Spp1 in TPO (red) and EV (blue) singlets (upper panel) and doublets (lower panel). (C) Volcano plot for differential gene expression of OsteoCAR singlet cells. Genes with logFC > 1 and adjusted p-value < 0.05 shown in orange; genes with p-value < 0.05 but low logFC in blue and others in grey.

**Supplementary figure 4.**
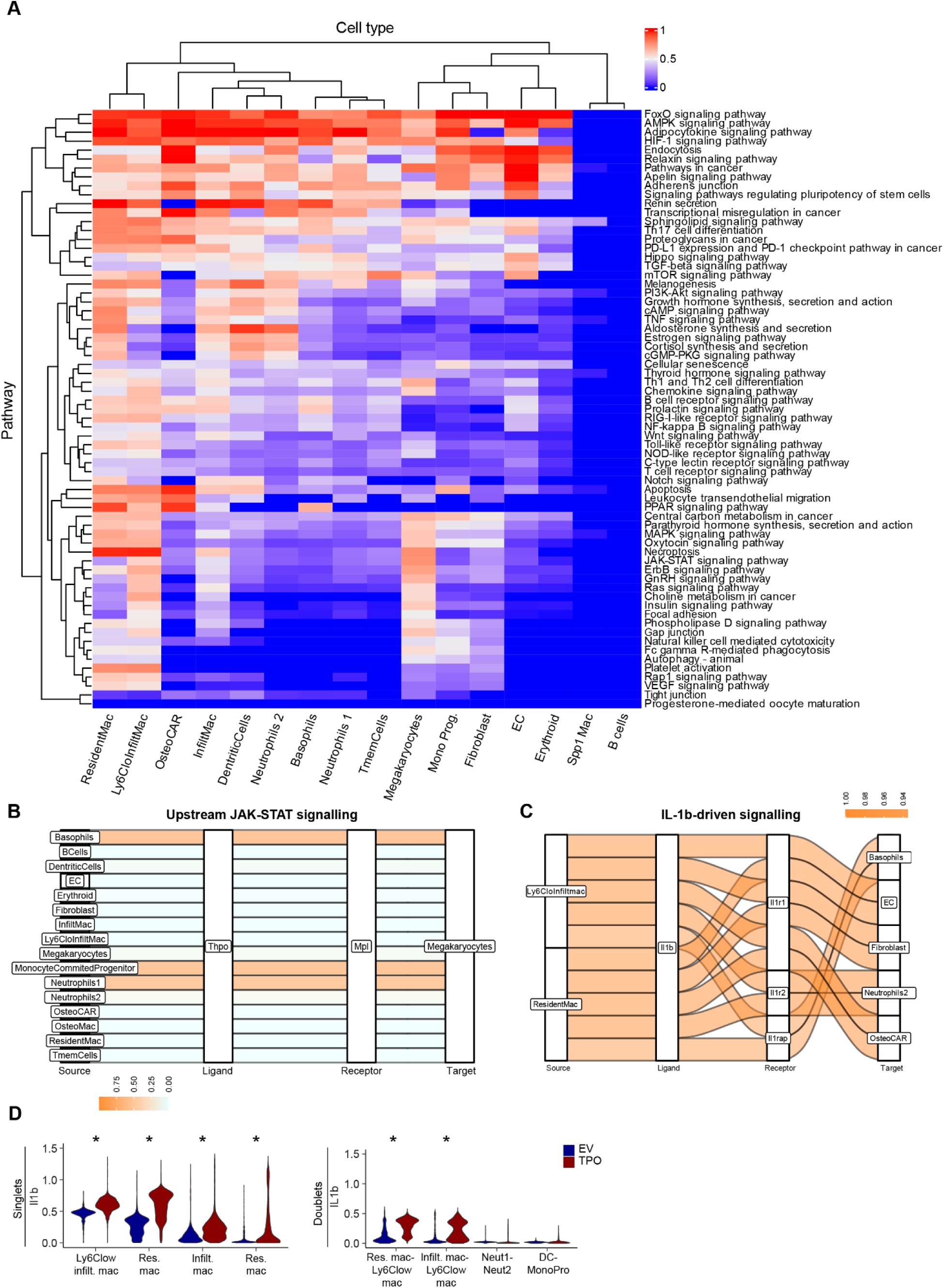
Macrophage-derived Spp1 promotes IL-1β–dependent inflammatory activation. (A) Heatmap of pathway score based on intracellular signaling across cell-types, highlighting differential transcriptional responses between TPO vs. EV. Higher scores indicate stronger evidence of differential pathway regulation at transcriptional level. **This indicates that the Jak/Stat pathway is particularly highly activated in Ly6C^low^ infiltrating macrophages and resident-like macrophages.** (B) Sankey plot showing Thpo-Mpl axis enriched in Megakaryocytes. Colors represent LR expression under TPO condition: orange indicates stronger signaling across cell-type pairs, while blue indicates weaker signaling. (C) Sankey plot showing the differential Il1b-associated interactions from Ly6C^low^ infiltrating macrophages and resident-like macrophages. Colors represent LR expression per interaction: orange indicates stronger signaling in disease, blue indicates weaker signaling in disease. (D) Violin plots showing expression of Il1b in TPO (red) and EV (blue) singlets (left panel) and doublets (right panel). Expression distributions difference significance measured via Wilcoxon rank sums tests.

**Supplementary figure 5.**
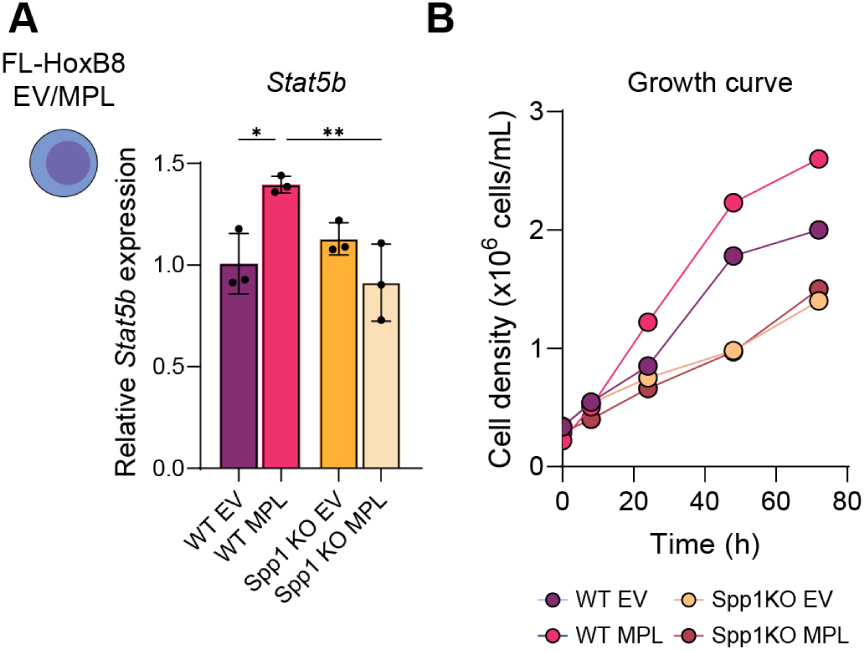
Deletion of Spp1 in hematopoietic cells carrying the MPLW515L mutation decreases JAK-STAT signaling and slows down clonal growth. (A) mRNA expression of Stat5b in WT or Spp1KO FL-HoxB8 cells (undifferentiated) carrying either the EV or MPLW515L mutation. n=3/group, P-value is based on a one-way ANOVA.(*) = p<0.05,(**)= p<0.01. (B) Growth curve of WT or Spp1KO FL-HoxB8 cells (undifferentiated) carrying either the EV or MPLW515L mutation. Y-axis is time in hours post-passaging.

**Supplementary figure 6.**
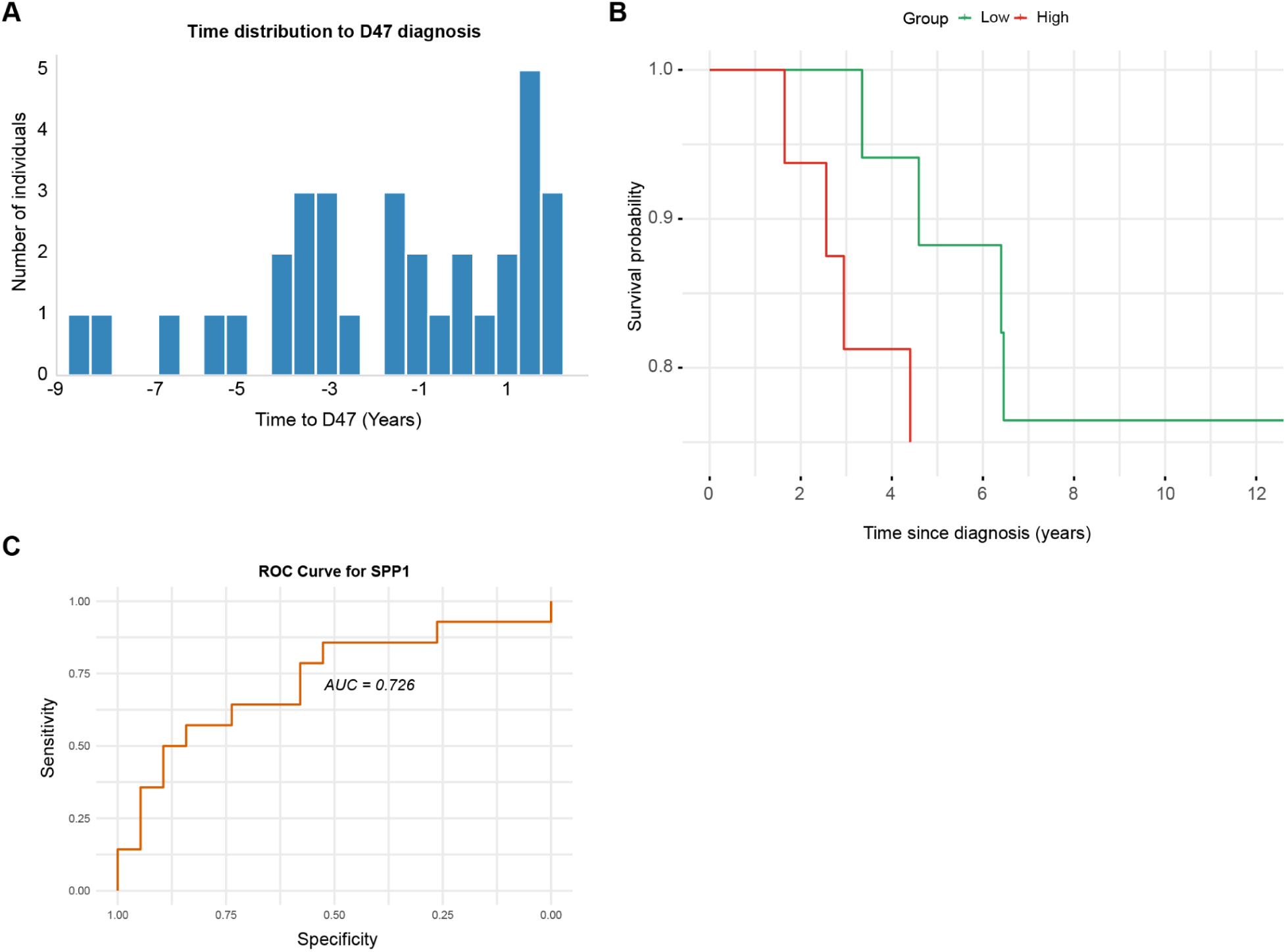
High circulating SPP1 correlates with inferior clinical outcomes in MPN. (A) Years to diagnosis (x-axis) vs. number of patients (y-axis) relative to the time of baseline assessment (ie. blood draw, year = 0) in the UK Biobank. (B) Kaplan–Meier survival curves of primary myelofibrosis patients stratified by low or high concentration of SPP1 concentration at baseline assessment. Differences between groups were evaluated using the log-rank test (C) ROC analysis for SPP1 showing the predictive ability of SPP1 to predict death in MPN patients

## Data and Software Availability

Nichesphere is provided as an open source python package at https://github.com/CostaLab/Nichesphere. Moreover, we provide tutorials on its use at https://nichesphere.readthedocs.io/en/latest/ and zenodo. We provide access to a Docker image, available at: https://gitlab.com/sysbiobig/ismb-eccb-2025-tutorial-vt3/container_registry. The Docker image comes preconfigured with all necessary libraries, tools, and software. Additionally, the repository at https://gitlab.com/sysbiobig/ismb-eccb-2025-tutorial-vt3 contains a summarized Nichesphere co-localization + communication analysis tutorial which was presented at the ISMB-ECCB 2025 conference. Raw singlet and multiplet cells data will be deposited in GEO, while pre-processed R and h5ad objects will be provided in Zenodo upon publication. The Zenodo reviewer access link to the data objects and Nichesphere tutorial is: https://zenodo.org/records/16781207?token=eyJhbGciOiJIUzUxMiJ9.eyJpZCI6IjQwZmU2NjdjLTU4ZWMtNDcyOS1iYzRkLWRiZjY4YzUyZDNiZSIsImRhdGEiOnt9LCJyYW5kb20iOiI4MTczYjQ4ODc0N2U5YjY1OGE3MDUyOWIwNzkxMDQwMSJ9.w1N59ymErmCjNs_nLKPL7-Tdoy_uS4fqZocDm7XvUcjRbgEQpjidtsEFUtiWvBGFwDPx7gLHkrFPxC_5Pm-bxQ

## Acknowledgments

We thank the team of the Laboratory Animal Science Center at Erasmus Medical Center, Rotterdam for excellent technical assistance. This research has been conducted using the UK Biobank Resource under Application Number 71300. UK biobank data was accessed by C.V.S and K.M.S. Copyright 2024, NHS England. Re-used with the permission of the NHS England and/or UK Biobank. All rights reserved. This work uses data provided by patients and collected by the NHS as part of their care and support.

H.G. was supported by a Gilead Research Scholar Award in Oncology/Hematology, a ZonMW VENI grant, an Erasmus Medical Center Fellowship and a KWF (Dutch Cancer Society) Exploration grant. G.C. was supported by a fellowship from the Fondazione Ing. Aldo Gini. R.K.S. is an Oncode Institute investigator and is supported by ERC grants (Rewind-MF ERC-CoG 101124542; deFIBER ERC-StG 757339 and PoC DeAlarmin) and a ZonMW VIDI grant. This work was in part supported by grants of the Deutsche Forschungsgemeinschaft (DFG) (German Research Foundation) to H.G (*417911533*), R.K. (KR 4073/9-1), R.K.S. (504777725; 417911533; 514007497) and I.C. (*417911533*). The project also received funding from the program “Netzwerke 2021”, an initiative of the Ministry of Culture and Science of the State of Northrhine Westphalia (CANTAR network). R.K., I.G.C., and R.K.S. are members of the E:MED Consortia Fibromap and the consortium CureFib funded by the German Ministry of Education and Science (BMBF). R.K.S and R.K. are co-founder and shareholder of Sequantrix GmbH. For the remaining authors, no relevant conflicts of interest were declared.

