## Supplemental file for "NicheSphere reveals Spp1⁺ macrophages as central hubs coordinating fibrotic remodeling in myeloproliferative neoplasms"

### Supplementary Material

### 1 NicheSphere Method Description

We describe here NicheSphere, which is a computational approach to combine spatial or physical interaction information with cell-cell communication to perform spatially constrained cell-cell communication analysis. NicheSphere has two main modules. The first one explores cell co-localization statistics to derive a cell-type-co-localization graph, and to delineate phenotype specific niches. The second module builds upon the cell-co-localization graph and niches to investigate cell-cell communication mechanisms associated to the conditions.

#### 1.1 Cell Co-localization

Cell type deconvolution methods produce a matrix $\mathbf{C} = \{\mathbf{c_{ik}}\}^{\mathbf{NxK}}$, where $c_{ik}$ is the probability of multiplet $i$ to be of cell type $k$, $N$ is the number of multiplets and $K$ is the number of cell types. Let $y_i \in \{1,...,k,...,K\}$ be a categorical variable indicating the cell type associated to a multiplet $i$. The cell type deconvolution (as computed above) provides us with the probability:

$$Pr(y_i = k) = c_{ik} \tag{1}$$

where $c_{ik}$ is the results of the previously defined cell type deconvolution. NicheSphere uses these results to estimate the probability of two cell types, defined as $\mathcal{X} \in \{1,...,K\}$ and $\mathcal{Y} \in \{1,..,K\}$, to be co-localized.

$$Pr(\mathcal{X} = k, \mathcal{Y} = j) = \frac{1}{N}\sum_{i=1}^{N} Pr(y_i = k)Pr(y_i = j) \tag{2}$$

The frequency of cell types in singlets is used to estimate background distribution for a pair of cells. That is for a particular cell type:

$$Pr(\mathcal{X}^s = k) = \frac{m_k}{M} \tag{3}$$

where $M$ represents the total number of singlets and $m_k$ the number of singlets associated to cell type $k$. Assuming the independence, the expected probability of seeing a pair of cell types in the singlet data is:

$$Pr(\mathcal{X}^s = k, \mathcal{Y}^s = j) = Pr(\mathcal{X}^s = k)Pr(\mathcal{Y}^s = j) \tag{4}$$

This can be used to define log odds scores by contrasting the probability of observing a pair in the multiplet vs singles:

$$L(\mathcal{X} = k, \mathcal{Y} = j) = log_2\left(\frac{Pr(\mathcal{X} = k, \mathcal{Y} = j)}{Pr(\mathcal{X}^s = k, \mathcal{Y}^s = j)}\right) \tag{5}$$

This provides us with a matrix $\mathbf{L} = \{\mathbf{l_{jk}}\}^{\mathbf{K \cdot K}}$ with the log odds scores for all the cell type pairs in a data set. Positive values indicate a higher chance to find a pair to be co-localized in the multiplet than in the singlet data, while negative values indicate a lower chance to find this pairs to be co-localized in the multiplet data than in the singlet data. All the above statistics can be estimated conditional on a condition, i.e. control $C$ or disease $D$, by only considering data (singlets and multiples) from that condition. These will define matrices $\mathbf{L^D}$ or $\mathbf{L^C}$.

Note also that the same principle can be used on an integrated analysis of spatial transcriptomics data as measured by spotted arrays. For this, NicheSphere requires a cell type deconvolution matrix $\mathbf{C}$ obtained by mapping a single cell data to the spatial spots using an state-of-art tool such as Cell2Location [8]. The statistics for singlets (Eq. 3) can be estimated from the reference single cell data accordingly.

##### *1.1.1 Statistical tests*

NicheSphere implements distinct statistical tests depending on the study design at hand. They include a test where samples are only measured in a single condition, two conditions without replicates; and finally two conditions with replicates. These are described below.

**Condition Specific and single sample cell pair enrichment test**

We next resort to empirical $p$-values to evaluate the significance of the co-occurrence statistics for a pair of cells for a particular study (condition specific or not). To formally assess the significance of the observed co-occurrence, we test:

$$\begin{cases} H_0 : \theta = \theta_0 & \text{(No significant co-occurrence)} \\ H_1 : \theta \neq \theta_0 & \text{(Significant co-occurrence)} \end{cases}$$

where $\theta$ represents the observed co-occurence probability and $\theta_0$ is the expected co-occurrence probability under the null model, derived from permutations of singlets. The alternative hypothesis posits that the observed co-occurrence deviates significantly from this null model, thus indicating a potentially meaningful association.

For this, we create random multiplets by sampling pairs of singlets data from the experiments to create multiplet data sets with $n$ observations. The true labels of the singlets indicate the composition of the random multiplet $C^r$. This is next used to compute a matrix $\mathbf{L^r}$ as defined in Eq. 5. This is repeated for $l$ times (default 10); and we only consider cell types with $L$ scores in the top (or bottom) 2.5% quartiles.

**Two conditions and single sample cell pair enrichment test**

For the comparative case with two conditions ($Z \in \{D, C\}$) each from a single sample, we can define:

$$L(\mathcal{X} = k, \mathcal{Y} = j) = log_2\left(\frac{Pr(\mathcal{X} = k, \mathcal{Y} = j|Z = D)/Pr(\mathcal{X}^s = k, \mathcal{Y}^s = j|Z = D)}{Pr(\mathcal{X} = k, \mathcal{Y} = j|Z = C)/Pr(\mathcal{X}^s = k, \mathcal{Y}^s = j|Z = C)}\right). \tag{6}$$

We use the same randomization and empirical p-value approach as described before to generate random scores; and a to obtain p-values.

**Two conditions and multiple samples cell pair enrichment test**

Here, we are interested in the case where we have several samples related to two conditions. For this, we define a vector $\mathcal{Z} = (z_1, ..., z_m)$, where $z_i \in \{D, C\}$ indicates if the sample is related to the disease ($D$) or control ($C$) group. For each sample $i$, we will have the probabilities of every pair of cell types $k$ and $j$ to be co-localized $Pr(\mathcal{X} = k, \mathcal{Y} = j)$ as defined in Eq. 2 For a pair of cells $k$ and $j$, we use a rank sum test [11] to compare the distribution of co-localization probabilities between two groups.

Formally, we test:

$$\begin{cases} H_0 : F_1(x) = F_2(x) & \text{(the two groups follow the same distribution)} \\ H_1 : F_1(x) \neq F_2(x) & \text{(the two groups differ in distribution)} \end{cases}$$

where $F_1$ and $F_2$ denote the cumulative distribution functions of co-localization probabilities in each group. The rank sum test evaluates whether the observed differences in ranks are consistent with the null hypothesis, under which any observed difference in co-localization probabilities between the two groups is attributed to random variation. The alternative hypothesis suggests that the two groups have systematically different distributions, indicating a condition-specific effect on co-localization.

We only consider cell-type pairs with significant changes (default $p$-value < 0.05). We use the W statistic (sum of the ranks in the disease condition) from the rank sum test as final weights in the cell-co-localization network.

#### 1.2 Cell co-localization graph

From the previous test results, we built a co-localization graph, where nodes represent cell types. Edges are included between cell types whenever their are detected to be significantly co-localized by the statistical test. The adjacency matrix for this undirected graph will be denoted as $\mathbf{A} = \{\mathbf{a_{jk}}\}^{\mathbf{K \cdot K}}$ with entries proportional to the Euclidean distance $E_{dist}$ as estimated on the co-localization scores $L$. More formally, this is defined by:

$$a_{jk} = 1 - \frac{1}{M} \cdot Euclidean(j,k) \tag{7}$$

where

$$Euclidean(j,k) = \sqrt{\sum_{i}^{k}(l_{ij} - l_{ik})^2} \tag{8}$$

and $M$ is the largest Euclidean distance value in the data. This formulation transform the Euclidean distance to a similarity values varying from 0 to 1.

We use this weighted graph to generated a graph embedding for visualization purposes using neato layout [6]. The same graph is used as input to detect niches via the community detection algorithm Louvain [1]. This provides a grouping of cell types into $o$ niches, as defined by the variable $c_k \in \{1,...,o\}$. This graph is also used to estimate model level properties such as betweeness scores [4], page rank [9] and signed node degree [5]. This is used to rank cell types regarding their connectivity.

### 2 Cell-Cell Communication test

#### 2.1 Cell-Cell communication graphs

Next, NicheSphere obtain a cell-cell communication graph by applying a ligand receptor algorithm as LIANA [2] from a singlet and condition specific scRNA-seq data. This is represented by a list of 5-tuples $X = \{x_1,...,x_n\}$ comprising $n$ significant LR interactions. A tuple is defined as $x_i = (s, g_{lig}, r, g_{rec}, e)$, where $s \in \{1,..,k\}$ is the sender cell type, $r \in \{1,..,k\}$ is the receiver cell type, $g_{lig}$ represents a ligand gene, $g_{rec}$ is a receptor gene and $e > 0$ is a positive weight of the LR interaction. The weight represents the ligand-receptor scores of the pair, such as the harmonic mean of the ligand and receptor genes as defined in CellPhoneDB [3]. In case of multiple conditions, this analysis is performed for every condition individually.

We obtain a directed cell-type graph by converting the 5-tuples using the following operation. For a given directet cell pair, the edges between $s$ to $r$ is represented by the following distribution:

$$w_{s,r} = \{e|(s, g'_{lig}, r, g'_{rec}, e) \in S\} \tag{9}$$

To contrast two cell-cell communication graphs $W^D$ and $W^C$, we use the rank sum test [11] as described above, to compare the distribution of a pair of cells ($W^D_{s,r}$ vs $W^C_{s,r}$).

Of note, we only consider cell type pairs present in the co-localization graph, that is $a_{sr} > 0$. The test provide us with a W statistic (sum of the ranks in the disease condition), which is used as weight in the directed cell type graphs for visualization purposes.

#### 2.2 Ligand-receptor mechanisms

To improve the interpretation of cell-cell communication results, NicheSphere uses an annotation of LR pairs $(g_{lig}, g_{rec})$ in mostly mutually exclusive mechanism categories $Q \in \{1, ..., q\}$ extracted from distinct databases as Matrisome [10] and Cytosig [7] . In total, ten mechanism categories are provided by NicheSphere by extracting ligands related to extra-cellular matrix (ECM) related mechanisms and immune cell recruitment. NicheSphere uses these ligands to find LR pairs associated with each mechanism category. See Table **??** for listing of process and number of LR interactions.

| **Process** | **Database** | **# ligands** | **#LR pairs** |
|---|---|---|---|
| Collagens | Matrisome | 45 | 321 |
| ECM affiliated | Matrisome | 129 | 143 |
| ECM glycoprots | Matrisome | 168 | 422 |
| ECM regulators | Matrisome | 204 | 195 |
| Proteoglycans | Matrisome | 36 | 43 |
| Secreted Factors | Matrisome | 265 | 1225 |
| Chemokine | Cytosig | 42 | 280 |
| Cytokine | Cytosig | 86 | 237 |
| Growth Factors | Cytosig | 130 | 1064 |
| Inhibitory | Cytosig | 36 | 49 |

#### 2.3 Mechanism and niche specific differential communication network

NicheSphere uses next these categories to define mechanism specific and niche specific cell-cell communication networks. In short, the sets $W_{s,r}$ can be filtered by only considering ligand-receptor pairs $(g_{lig}, g_{rec})$ from a particular mechanism $Q$, i.e. $W^Q_{s,r}$. This filtered LR sets can be then used in the rank sums tests as previously defined.

Moreover, NicheSphere assumes that cell-cell communication are niche specific. Therefore, for a pair of niches $i$ and $j$, such that $1 \leq i < j \leq o$, this can be done by the union of the $W_{s,r}$ pairs contained in the niches. This allow us to perform a condition and niche specific test.
